# A comparative framework for neocortical sulcal anatomy in pinnipeds

**DOI:** 10.64898/2026.09.17.752275

**Authors:** Magdalena Boch, Kamilla Avelino-de-Souza, Sara Binder, Nina Patzke, R. Austin Benn, Heitor Mynssen, Khallil Taverna Chaim, Khin Khin Tha, Bridget Wicinski, Cheuk Y. Tang, Paul R. Manger, Andrew A. Rouse, Patrick R. Hof, Peter F. Cook, Rogier B. Mars

**Author notes:** shared authorship (Magdalena Boch, Kamilla Avelino-de-Souza). Corresponding authors: Magdalena Boch; Kamilla Avelino-de-Souza.

## Abstract

The transition of pinnipeds from terrestrial ancestors into marine environments provides a unique opportunity to examine how cortical organisation has changed with adaptation to life in water. Here, we examined neocortical sulcal anatomy in nine species representing all three extant pinniped families using cortical surfaces reconstructed from post-mortem MRI. Building on a standardised framework previously established for land-dwelling (fissiped) carnivorans, we developed identification criteria for major pinniped sulci and compared their configurations with those observed across fissipeds. Major sulci were broadly conserved, but pinnipeds exhibited characteristic modifications, including a complex, highly branched pseudosylvian configuration and distinct frontoparietal sulcal patterns, with further family-level variation in both territories. This observed variation in sulcal patterns may reflect pronounced differences in locomotor, tactile, and vocal specialisations, generating hypotheses about how cortical organisation may vary with adaptation to marine environments. Together, these findings establish a common anatomical framework across Carnivora for investigating how cortical organisation diversified in relation to behaviour and ecology.

## 1. Introduction

The evolutionary transition from terrestrial ancestors into marine environments is rare among mammals. Within the mammalian order Carnivora, pinnipeds (from *pinna*, fin, and *pes*, foot) represent the most species-diverse clade to have undergone this transition [1,2]. They form a monophyletic lineage within Caniformia comprising three extant families: Odobenidae (walruses), Otariidae (fur seals and sea lions), and Phocidae (true seals). Compared with the remaining primarily land-dwelling carnivorans, often collectively referred to as fissipeds, pinnipeds comprise relatively few extant species yet exhibit pronounced morphological adaptations to marine environments [3,4]. They therefore provide a unique comparative opportunity to examine how the carnivoran cortical *Bauplan* has been modified following a major ecological transition.

More broadly, the order Carnivora provides a powerful natural experiment for studying brain evolution. With representatives occupying terrestrial, arboreal, fossorial, and semi-aquatic niches, carnivorans exhibit striking ecological and behavioural diversity despite sharing a relatively recent common ancestor approximately 55 million years ago [5]. Comparative work across the order has revealed variation in encephalization, brain shape, and neuroanatomical organisation associated with ecological and behavioural specialisations [6–12], including a more highly gyrencephalic cortex in pinnipeds compared to fissiped carnivores and primates of similar brain size [13–15]. However, our understanding of carnivoran brain diversity remains incomplete and unevenly distributed across lineages, with many aspects of pinniped brain organisation still poorly understood.

This gap is particularly apparent at the level of cortical organisation. While global measures such as brain size and shape provide important insights into broad evolutionary patterns, understanding how brains differ across species also requires examining regional aspects of cortical organisation [16,17]. Yet comparatively little is known about how the pinniped neocortex is organised, how its morphology varies across the three extant families, and how it relates to that of fissipeds. Establishing corresponding cortical features across these species is therefore an important first step towards investigating which aspects of cortical organisation reflect their shared carnivoran ancestry and which may have been modified during adaptation to marine environments.

Neocortical sulci provide clear macroanatomical landmarks that can be used to establish such a comparative reference framework across species and investigate lineage-specific modifications in cortical folding patterns [18–20]. In a recent study, we established a standardised framework for identifying and comparing major sulci across fissipeds [21], providing a reference for systematic interspecies comparisons. For pinnipeds, however, detailed comparative descriptions of neocortical sulcal anatomy remain scarce. Existing accounts typically focus on a small number or single species [22–24], selected cortical regions [24–28], or endocranial casts [29,30], and often employ inconsistent terminology. As a result, corresponding sulci across pinniped families and between pinnipeds and fissipeds remain insufficiently resolved, and a standardised framework for identifying and comparing pinniped sulci is currently lacking.

Recent advances in comparative neuroimaging and collaborative brain collections now make it possible to address this gap by enabling systematic comparisons of neocortical organisation across rarely studied, phylogenetically diverse species [31–34]. Here, we use cortical surfaces reconstructed from post-mortem MRI to examine neocortical sulcal anatomy across representatives of all three pinniped families, building on our previously established framework for fissiped carnivorans.

Our aims are threefold. First, we develop standardised protocols for identifying major pinniped sulci. Second, we provide a comparative description of sulcal organisation across families. Third, we assess which aspects of pinniped sulcal anatomy are conserved relative to fissipeds and which represent lineage-specific modifications. Together, these comparisons establish a common anatomical reference across Carnivora and lay the foundation for future work into the relationship between cortical organisation and ecological specialisations.

## 2. Results & Discussion

We labelled the neocortical sulci of nine pinniped species representing all three extant families (**Figure 1**) using standardised identification criteria originally developed for fissipeds [21] and adapted these to pinniped sulcal anatomy. Additional individuals were included for several species to assess intraspecific variation (**Supplementary Table 1**).

**Figure 1.**
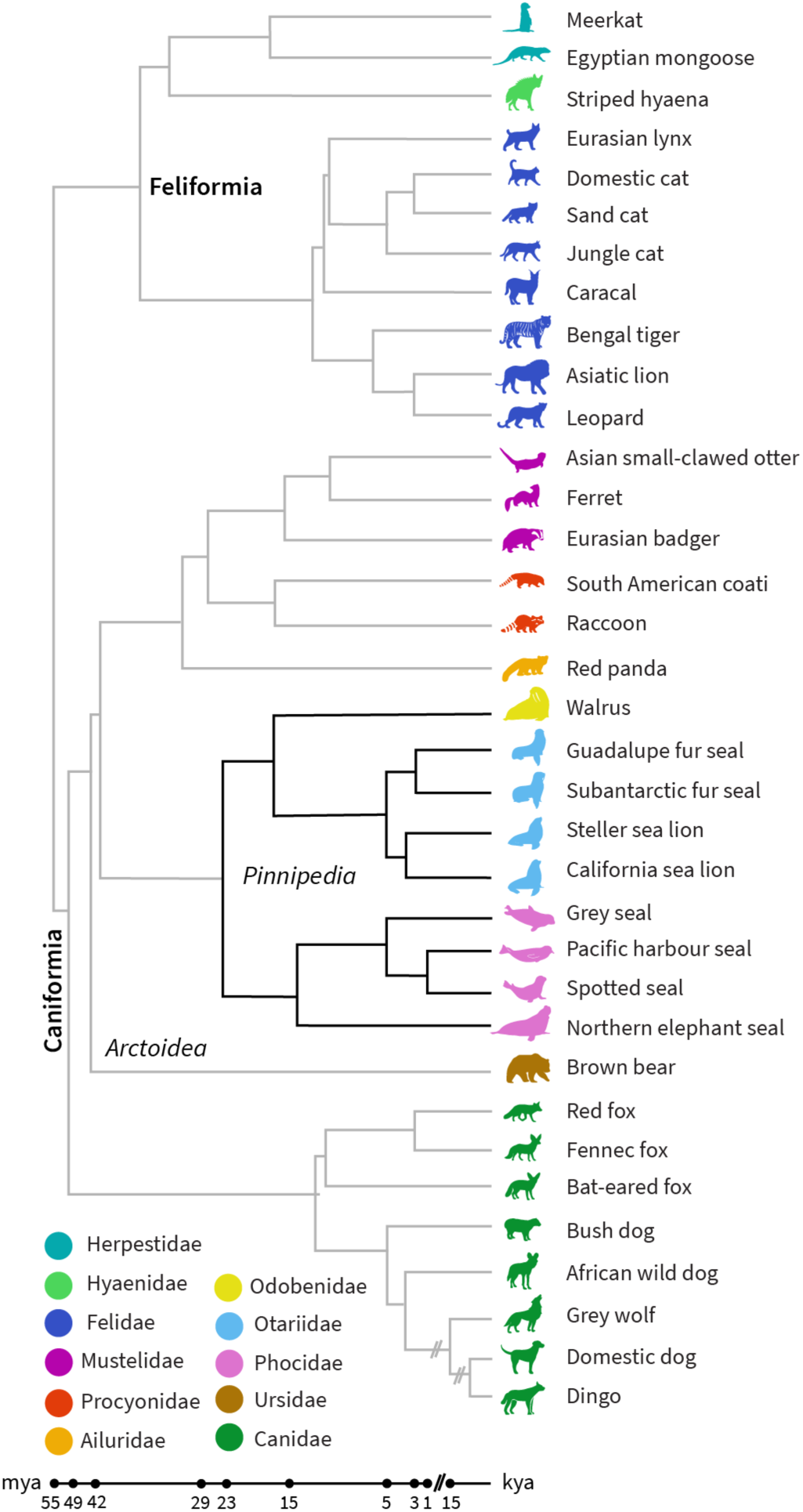
Phylogeny of carnivoran sample. The sample comprises nine pinniped species representing all three extant families (see **Table 1**). Branches representing previously described fissiped carnivorans [21] are shown in grey for comparison. Phylogenetic relationships were derived from [5,35–37]. k/mya, thousand/million years ago.

Overall, pinniped brains appeared more convoluted, with deeper folds and a greater number of secondary sulcal branches, than fissipeds with comparable brain mass (15,21). Larger pinniped brains also tended to exhibit more pronounced sulcal complexity than smaller brains, consistent with previous observations across carnivorans [15,21]. We observed no pronounced hemispheric differences in major sulcal patterns. Below, we describe the morphology and identification of each major sulcus and variation across species.

### 2.1. Identification of major sulci in the carnivoran neocortex

#### Occipitotemporal region

##### Pseudo-sylvian (or sylvian) fissure or complex

The pseudosylvian fissure originates at the ventral border of the neocortex (**Figure 2**, cyan) and was present in all examined pinniped brains. It extends dorsocaudally and, as in other arctoid carnivorans, appears more elongated than in other carnivoran clades [cf. 21]. In contrast to fissipeds, however, the fissure exhibits a reduced caudal curvature, and in some species, including the Guadalupe fur seal, it is oriented nearly perpendicular to the ventral surface. In the pinnipeds studied, the pseudosylvian fissure displayed additional secondary branches, resulting in a more elaborate sulcal configuration, consistent with earlier descriptions in pinniped brains [22]. Given this variability in shape and branching pattern, we refer to this structure as a pseudosylvian sulcal complex rather than a single fissure, encompassing both the fissure proper and associated sulcal branches.

**Figure 2.**
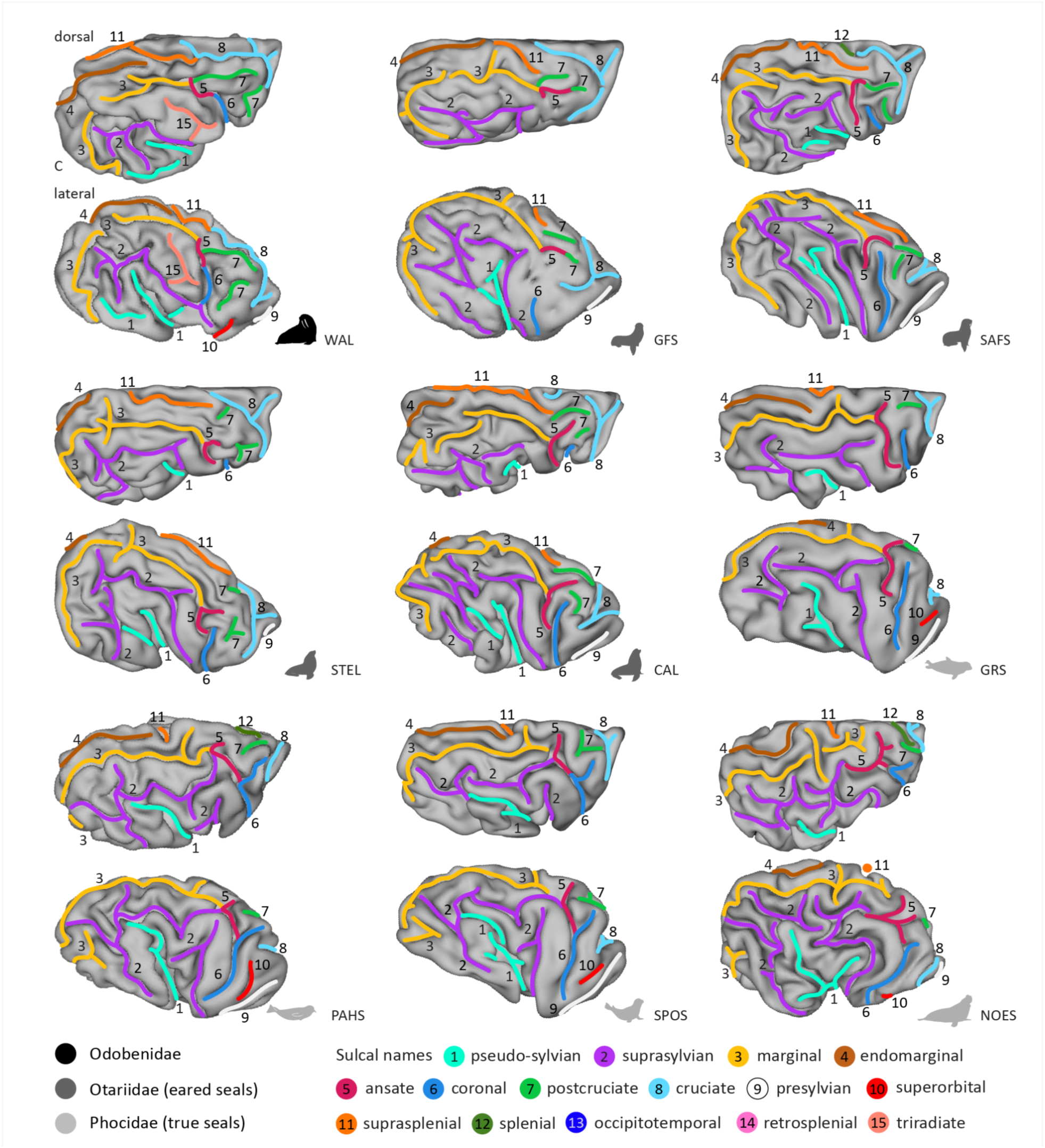
Lateral and dorsal views reveal varying sulcal complexity in parietal and occipitotemporal cortex between clades, as well as medial wall sulci crossing the midline. All species exhibit complex sulcal configurations associated with the pseudosylvian fissure (1, cyan), here termed the pseudosylvian complex. The suprasplenial sulcus (11, orange) crosses the midline onto the dorsal surface in all pinnipeds and extends further in otariids than phocids. Phocids generally exhibit a more rostrally extended endomarginal sulcus (4, brown) and a detached rostral suprasylvian sulcus (2, purple), whereas odobenid and phocids species exhibit a superorbital sulcus (10, red); see also **Supplementary Figure S1** for additional individuals. Surfaces in all figures are sorted by pinniped families, indicated by icon colours: black, Odobenidae; dark grey, Ottariidae; light grey, Phocidae. CAL, California sea lion; GFS, Guadalupe fur seal; GRS, grey fur seal; NOES, Northern elephant seal; PAHS, Pacific harbour seal; SAFS, subantarctic fur seal; SPOS, spotted seal; STEL, steller sea lion; WAL, walrus.

In all species examined, in all individuals (including the additional individuals described in the Supplementary Material; see **Supplementary Figure 1**), we observed an additional sulcal extension at the dorsal end of the pseudosylvian fissure. This dorsal extension was consistently more elongated in phocids than in otariids or odobenids, with the dorsal sulcus terminating closely to, or even merging with, the mid suprasylvian sulcus. A comparable configuration was also observed for a sulcus arising from the mid portion of the pseudosylvian fissure and bending caudally, which appeared more elongated in the majority of otariids compared to phocids, terminating closely or merging with the caudal suprasylvian sulcus. Although these structures do not constitute fissures in the strict sense, as they do not overlie non-neocortical tissue, they are, in the majority of species, continuous with the pseudosylvian fissure. We therefore include them as components of the pseudosylvian sulcal complex.

With few exceptions, the dorsal branch of the pseudosylvian sulcal complex exhibited a caudal bend. In the spotted seal, this bend merged with the caudal part of the mid suprasylvian sulcus (purple; **Figure 2, Supplementary Figure S1**). This pattern was present in all hemispheres, except for the left hemisphere of the first individual. In the subantarctic fur seal, this branch instead bent anteriorly, whereas in the Guadalupe fur seal, it extended in both anterior and caudal directions. In northern elephant seals, the dorsal extension bifurcated into a characteristic Y-shape, with the caudally directed branch being more elongated. This bifurcation is not clearly visible in the lateral projection shown in **Figure 2**, as it lies largely hidden beneath the ectosylvian gyrus (**Supplementary Figure S3**). In the first individual, the caudal sulcus merged with the posterior part of the mid-suprasylvian sulcus; in the second, it terminated close to the sulcus.

In otariids and odobenids, the sulcal branch arising from the middle portion of the pseudosylvian fissure differed from its phocid homologue. The greater difference in depth between the fissure proper and the adjoining sulcus gave the latter a more detached appearance. In addition, this sulcal branch was often more elongated. In all otariids, except for the subantarctic fur seal and the right hemisphere of the third California sea lion, the caudal sulcus superficially merged with the mid or caudal suprasylvian sulcus. In the walrus, however, the caudal branch did not merge with the suprasylvian sulcus but instead terminated in proximity to the posterior marginal sulcus. Moreover, a second, less pronounced sulcus arose anteriorly from the mid portion of the pseudosylvian fissure, giving the overall configuration a more bifurcated appearance in this species.

##### Ectosylvian sulcus

In fissipeds, when present, the ectosylvian sulcus forms a continuous arch in canids, is interrupted at its midpoint in felids, and in herpestids and hyenids is represented only by a posterior sulcus, with no anterior component [21]. This sulcus has not been described in land-dwelling arctoid species, nor in previous anatomical descriptions of pinniped brains [e.g., 22,28,38].

In the pinnipeds examined here, we did not observe a sulcus closely resembling the fissiped ectosylvian sulcus in either overall shape or continuity. However, the caudally directed sulcal branch arising from the mid portion of the pseudosylvian fissure (described above) represents a candidate structure for a potential posterior ectosylvian sulcus, based on its position and trajectory relative to adjacent suprasylvian sulci. Determining whether this sulcus is homologous to the fissiped ectosylvian sulcus will require additional information on the cytoarchitecture and functional organisation of this cortical region. For this reason, we currently describe this sulcal branch as part of the pseudosylvian sulcal complex rather than terming it ectosylvian sulcus.

##### Suprasylvian (or supersylvian) sulcus

The suprasylvian sulcus is present in all specimens and, as in fissiped arctoids, forms the first arc around the pseudosylvian fissure (purple, **Figure 2**). However, unlike the typically continuous configuration observed in many land-dwelling arctoids, the sulcus exhibits prominent radial branches and is fragmented to varying degrees, with rostral, middle, and caudal components that may appear fully separated, partially continuous, or only locally detached depending on the species or lineage. The rostral component is also frequently obscured from lateral view by the rostral suprasylvian gyrus (i.e., the gyrus between the rostral suprasylvian and coronal sulcus; also called the coronal gyrus), likely contributing to its inconsistent reporting or misidentification in previous descriptions of pinniped sulcal anatomy [e.g., 24,38]. For illustration purposes, we therefore traced the ventral border of this gyrus in **Figure 2** to indicate the approximate position of the rostral suprasylvian sulcus (**Supplementary Figure S3** illustrates the hidden trajectory of the sulcus in example species). While this sulcus has been identified in previous studies [22,26], Fish [22] described the rostral portion as the ‘pre-supersylvian’ sulcus and distinguished it from a ‘post-supersylvian’ component.

In phocids and otariids, the middle portion of the suprasylvian sulcus either merges with or terminates in close proximity to the ansate (berry) sulcus, giving the appearance of a continuous suprasylvian arc (see, for instance, the Pacific harbour seal). In combination with the frequent detachment and concealment of the rostral suprasylvian sulcus, particularly in phocids, this configuration likely underlies previous reports in which the coronal sulcus has been identified as the rostral suprasylvian sulcus [24,38].

Within otariids (dark grey icons; **Figure 2**), the suprasylvian sulcus was divided at its mid-portion in the subantarctic fur seal and California sea lion. In the second and third California sea lion individuals, the caudal end was also detached (left hemisphere in the second individual; bilaterally in the third; **Supplementary Figure 1**). The Guadalupe fur seal had both a mid-portion split and a detached caudal suprasylvian sulcus. The Steller sea lion exhibited a detached caudal suprasylvian sulcus in all but the right hemisphere of the third individual, in which the rostral portion was detached instead.

Within phocids (light grey icons), the rostral suprasylvian sulcus was fully detached from the middle portion in the Pacific harbour seal and spotted seal, and only superficially connected in the grey seal. In the Northern elephant seal, the rostral portion remained continuous with the middle suprasylvian sulcus. In most phocids, the middle suprasylvian sulcus either fully or superficially merged with the ansate sulcus. In the grey seal and in the left hemisphere of the Northern elephant seal, however, the middle portion terminated in proximity to the ansate sulcus.

In odobenid (black icon), the suprasylvian sulcus of the walrus was not divided into multiple portions, and the middle suprasylvian sulcus did not extend or branch toward the ansate sulcus.

##### Ectomarginal (or ectolateral) sulcus

In canids, the ectomarginal sulcus forms an additional half-arc between the marginal and suprasylvian sulci. We did not observe this configuration in any species in our sample. Its absence in pinnipeds, and more broadly in other fissiped species, suggests that the ectomarginal sulcus represents a canid-specific specialisation.

##### Marginal (or lateral) sulcus

The marginal sulcus is an elongated sulcus running parallel to the midline (yellow, **Figure 2**) and merging with, or terminating in close proximity to, the perpendicularly oriented ansate sulcus (berry) at its rostral portion. In pinnipeds, the caudal portion extends ventrally, roughly following the trajectory of the suprasylvian sulcus (purple, **Figures 2-3**). Together, the marginal, ansate, and coronal sulci form an incomplete arc curving around the suprasylvian sulcus, resembling the configuration previously described for the brown bear.

**Figure 3.**
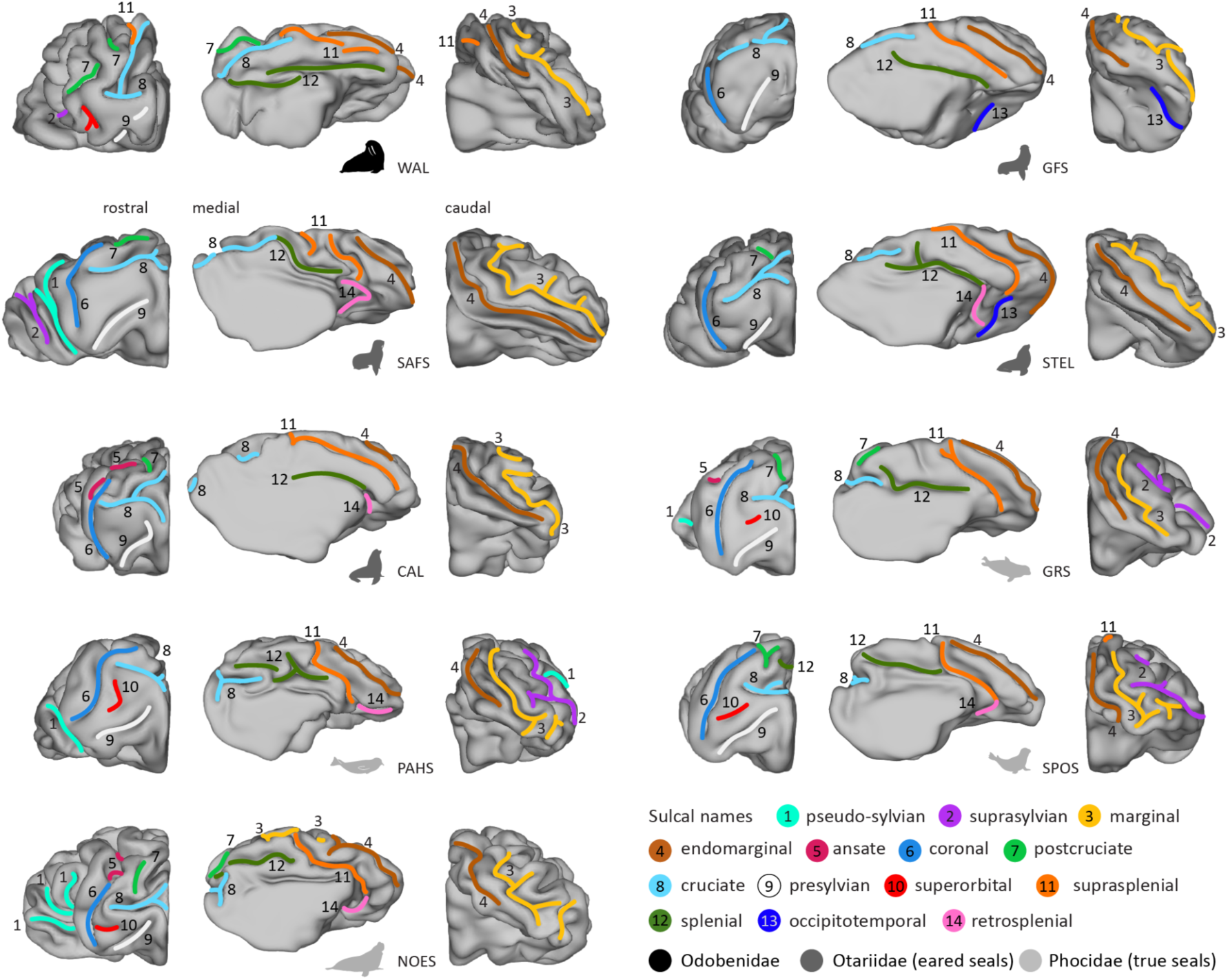
Rostral, caudal and medial views show lineage-specific parietal sulcal patterns and dorsal sulci extending onto the medial wall. Odobenids and phocids exhibit a superorbital sulcus (10, red) ventral to the cruciate sulcus (8, light blue). In all species, the endomarginal sulcus (4, brown) extends along the medial surface, while the splenial (12, dark green) and cruciate sulci generally terminate in close proximity rather than merging. See **Supplementary Figure S2** for additional individuals. Black, Odobenidae; dark grey, Otariidae; light grey, Phocidae. CAL, California sea lion; GFS, Guadalupe fur seal; GRS, grey fur seal; NOES, Northern elephant seal; PAHS, Pacific harbour seal; SAFS, subantarctic fur seal; SPOS, spotted seal; STEL, steller sea lion; WAL, walrus.

We did not observe marked interspecific variation in the overall configuration of the marginal sulcus, although its continuity and local morphology varied. In most species, a segment of the caudal portion was detached. In otariids (e.g., Subantarctic fur seal) and odobenids, this detachment typically occurred at a more dorsal point, where the sulcus began to curve ventrally, whereas in phocids exhibiting a detached caudal portion (e.g., Pacific harbour seal), the detachment occurred closer to its caudoventral termination. The caudal portion also tended to exhibit more secondary branching in phocids, resulting in a less continuous and more irregular trajectory than the comparatively smooth curvature observed in otariids and odobenids. Exceptions within Otariidae were the second California sea lion, which exhibited a continuous sulcus and the two additional Steller sea lion individuals, in which the sulcus exhibited a secondary split at its ventral bend (see **Supplementary Figures 1-2**). Within Phocidae, the grey seal and the second spotted seal likewise had a continuous sulcus.

##### Endomarginal (or entolateral, endolateral) sulcus

The endomarginal sulcus runs roughly parallel to the marginal sulcus along its medial aspect (brown, **Figure 2-3**). In fissiped arctoids, where present, it is typically located between the marginal sulcus and the longitudinal fissure and is comparatively short and shallow. In pinnipeds, the sulcus is more pronounced, both in depth and extent. It extends further caudally, often curving around the ventral bend of the caudal marginal sulcus, while its rostral portion continues onto the medial wall, where it runs approximately parallel to the suprasplenial sulcus (orange, **Figure 3**).

Across pinniped species, the endomarginal sulcus exhibited variation in its rostral and caudal extent. In odobenids and otariids, the caudal end was particularly elongated and extended far into the occipital region. In these species, the sulcus’ rostral extent was more limited, coinciding with the suprasplenial sulcus crossing the midline and extending anteriorly onto the dorsal surface.

In phocids, the endomarginal sulcus exhibited the opposite pattern, extending further rostrally while remaining more restricted caudally. The suprasplenial sulcus again defined the anterior boundary, but unlike in otariids and odobenids, it crossed the midline more anteriorly, to a lesser extent, and more perpendicularly to the endomarginal sulcus.

#### Frontoparietal region

##### Ansate and coronal sulcus

The rostral end of the marginal sulcus coincides with a complexly organised region comprising two distinct sulci. The first sulcus runs dorsomedially towards the medial longitudinal fissure and is termed the ansate sulcus (berry, **Figure 2**). The second sulcus continues rostrally on the lateral surface, curving ventrally, and is termed the coronal sulcus (medium blue, **Figure 2**). Across carnivorans, the coronal sulcus varies in its orientation and degree of ventral curvature relative to the marginal and suprasylvian sulci. In fissipeds, it typically forms a rostral extension of the marginal sulcus, exhibiting only a slight ventral curvature around the cruciate sulcus. In pinnipeds, the coronal sulcus is more strongly curved ventrally, oriented approximately perpendicular to the midline and aligning with the trajectory of the rostral suprasylvian sulcus. Together with the marginal and ansate sulci, this configuration forms an additional, partially detached arc around the suprasylvian sulcus. As noted above for the suprasylvian sulcus, the rostral suprasylvian, coronal, and ansate sulci inconsistencies in labeling have been noted and discussed before [22,28].

In phocids and most otariid species in our sample, the rostral marginal sulcus merged with the ansate sulcus near its middle portion or terminated in close proximity. In these cases, the ansate sulcus was oriented approximately perpendicular to the marginal sulcus and often exhibited a slight caudal bend around its middle portion. An exception to this pattern was the Guadalupe fur seal, in which the ansate sulcus appeared to extend primarily dorsally, with the marginal sulcus merging at its ventral termination; however, interpretation of this region was limited by the reduced quality of the reconstruction. The rostral marginal sulcus of the walrus merged with the dorsal tip of the ansate sulcus, and the sulcus exhibited a less pronounced bend. Within phocids in our sample, a branch of the middle suprasylvian sulcus (superficially) merged with or terminated near the ansate sulcus, which may contribute to the misidentification of the coronal and suprasylvian sulci noted above.

The coronal sulcus was consistently positioned more caudally than the cruciate sulcus and extended largely in a dorsoventral direction, following a relatively straight course. The relationship between the coronal and ansate sulci varied across otariid and phocid species and individuals (**Supplementary Figure S1**), with the two sulci appearing either superficially continuous, such as in the Stellar sea lion and northern elephant seal, or remaining distinct but terminating in close proximity, such as in the grey seal and the Subantarctic fur seal. In odobenids, the ansate sulcus of the walrus formed the intersection with the marginal sulcus, while the coronal sulcus merged with the rostral portion of the suprasylvian sulcus.

##### Cruciate (or medial cruciate) sulcus

The cruciate sulcus is positioned approximately perpendicular to the median longitudinal fissure on the dorso-frontal convexity of the brain. In pinnipeds, it occupies a comparatively more rostral position, closer to the frontal pole than in fissipeds (light blue, **Figures 2-3**), consistent with recent findings that pinniped brains are less rostrocaudally elongated when viewed from the dorsal perspective than those of fissipeds [9]. On the medial wall, the sulcus typically terminates close to the splenial sulcus. In pinnipeds, the cruciate sulcus exhibits a more complex morphology than in many fissipeds and frequently forms a Y- or T-shaped configuration. This pattern arises from a principal branch extending from the midline and curving ventrally onto the lateral surface, which bifurcates dorsally into a rostrally directed branch that curves towards the median longitudinal fissure. The laterally directed component has occasionally been termed the medial cruciate sulcus [29], but both components form a continuous, branching sulcal configuration.

The curvature and length of the cruciate sulcus’ lateral branch varied. In phocids, it extended more perpendicularly from the longitudinal fissure and did not curve as far ventrolaterally, rendering it less visible from a lateral perspective than in otariids and odobenid, as exemplified by the grey seal and Steller sea lion (lateral view, **Figure 2**). In otariids and odobenids, the sulcus also originated more caudally than in phocids (dorsal view, **Figure 2**).

##### Postcruciate and triradiate sulcus

The postcruciate sulcus (light green, **Figures 2-3**) is located between the ansate, coronal and cruciate sulci. In the walrus, an additional sulcus with a triradiate configuration (salmon, **Figure 2**) may be present between the ansate/coronal region and the rostral suprasylvian sulcus. While this configuration resembles the shape of the triradiate sulcus described in raccoons [39], the term is used here descriptively and does not imply homology.

In the odobenid and otariid species examined, which also exhibit an elongated cruciate sulcus with a more caudal origin (as described above), the cruciate sulcus often gives rise to an additional branch, and a secondary sulcus is present dorsal to the primary cruciate sulcus. This increased complexity may be described as a “postcruciate complex”, following its use in Procyonidae to denote a similarly complex configuration [21,39,40]. In some individuals, the cruciate sulcus appeared to merge with the ansate sulcus (berry, **Figure 2** and **Supplementary Figure S1**), as observed in the walrus, Guadalupe fur seal, and the third California sea lion (right hemisphere). However, in the former two species, this configuration should be interpreted with caution due to the limited quality of the underlying brain samples.

In the walrus, the additional triradiate sulcus exhibited a bifurcation at its rostroventral end, but this observation is based on a single hemisphere of limited quality and should be interpreted with caution. Previous accounts did not describe this sulcus, although digital endocasts suggest additional indentations in this region [29].

##### Presylvian sulcus

Rostral to the pseudo-sylvian fissure, the presylvian sulcus originates from, or close to, the rhinal fissure and extends rostrally with a gentle dorsal curvature (white, **Figure 3**). Across carnivorans, its position varies: in fissiped species, the sulcus typically terminates on the dorsal surface, whereas in pinnipeds it is consistently shifted ventrally. As a result, in pinnipeds the presylvian sulcus is less apparent in dorsal view but can be readily observed from a rostral perspective.

We did not observe major variation across species or individuals, except for a frequent detachment of the presylvian sulcus from the rhinal fissure (see also **Supplementary Figure S2**).

##### Proreal and diagonal sulcus

A proreal or diagonal sulcus was not observed in any of the pinniped species examined. This rostral frontal proreal sulcus is typically present in wolf-like canids, and the lateral frontal diagonal sulcus in felids and hyaenids [21,41,42]. Rioch [24] describes a proreal sulcus in the harbour seal, although this was not identified in other anatomical descriptions of pinniped brains [22,28,38], and it was not observed in our specimens.

##### Superorbital sulcus

The superorbital sulcus is located in the frontal lobe (red, **Figures 2-3**), positioned ventral to the cruciate sulcus (light blue) and dorsal to the presylvian sulcus (white). It occupies a more lateral and ventral position than the proreal sulcus of canids, which is typically situated rostral to the presylvian sulcus on the dorsal surface. It is also distinct from the diagonal sulcus of felids and hyenids, which lies caudal to the coronal sulcus, whereas the superorbital sulcus is positioned rostral to this region and corresponds to the “superorbital fissure” described by Fish [22].

We observed this sulcus in all phocid species examined, with variation in curvature and shape across species and individuals. For example, in the Pacific harbour seal, the sulcus exhibited a marked dorsal bend, whereas in the Northern elephant seal it showed only a gentle dorsal curvature (**Figure 3**). In the second Northern elephant seal (**Supplementary Figure S2**), the sulcus instead displayed a slight ventral curvature. In odobenids, the walrus also exhibited a superorbital sulcus, including a small secondary branch.

In otariids, the superorbital sulcus was not consistently identifiable. In some individuals, shallow or fragmentary indentations were observed in a comparable topological position, but these lacked the continuity and depth observed in phocids and odobenids to be confidently labelled as a distinct sulcus. Fish [22] described a corresponding superorbital sulcus in otariids, including the fur seal and the California sea lion. Our observations suggest that the sulcus may be present in otariids, but is less consistently expressed and comparatively shallow, and therefore not reliably identifiable on cortical reconstructions.

#### Medial wall region

##### Splenial and retrosplenial sulcus

The dorsal longitudinal splenial sulcus (dark green, **Figure 3**) runs rostrocaudally in close proximity to the splenium of the corpus callosum. At its rostral end, it approaches the cruciate (light blue) and postcruciate (light green) sulci and, in some cases, merges with either of them.

The sulcus curving ventrally around the posterior end of the corpus callosum is referred to as the retrosplenial sulcus (rosé). In pinnipeds, the retrosplenial sulcus either merges with the caudal end of the splenial or suprasplenial (orange) sulcus or remains detached. In fissipeds, the cruciate, splenial, and retrosplenial sulci are typically continuous [21].

Both sulci showed variation across individuals (**Supplementary Figure S2**). The splenial sulcus was present in all species examined, appearing either continuous or segmented and sometimes exhibiting a secondary dorsal branch at its rostral end. The retrosplenial sulcus additionally varied in its presence and could not be identified in the walrus. Where present, it merged with or terminated near the caudal splenial or suprasplenial sulcus.

##### Suprasplenial and occipitotemporal sulcus

The suprasplenial sulcus (orange, **Figures 2-3**) is located dorsal to the splenial sulcus and runs approximately parallel to it at its caudal extent. In fissipeds, the sulcus typically maintains this trajectory, curving around the posterior end of the splenial sulcus and remaining confined to the medial wall. In pinnipeds, the sulcus curves dorsally, crossing the longitudinal fissure and extending onto the dorsal surface, where it becomes visible rostral to the endomarginal sulcus (brown, **Figure 3**). The occipitotemporal sulcus (dark blue) curves around the retrosplenial sulcus.

Across pinniped lineages, the suprasplenial sulcus varied primarily in the degree of its dorsal extension, i.e., its extent onto the dorsal surface. In the odobenid and otariids examined, the sulcus emerged caudally and extended along the dorsal surface approximately parallel to the marginal sulcus, terminating near, or merging with, the postcruciate sulcus (light green, **Figure 3**. In phocids, by contrast, the sulcus reached the dorsal surface more rostrally, near the dorsal end of the marginal sulcus, and remained comparatively short, such that it was typically not visible from the lateral surface, as seen in the Northern elephant seal. These differences co-occurred with variation in the rostrocaudal extent of the endomarginal sulcus, which extended further rostrally in phocids and remained more restricted in otariids and the walrus.

The occipitotemporal sulcus could not be clearly identified in the phocids and the walrus, being only variably present in otariids. Where identifiable, it terminated near the caudal end of the suprasplenial sulcus but remained detached from it.

### 2.2. Lineage-specific observations and potential relationship to function

Our sulcal framework revealed regional variation in cortical organisation across pinniped lineages. Several of these differences occur within or adjacent to known sensory and motor cortical territories and coincide with variation in locomotor and sensory specialisations (**Figures 5-6**). Although the mechanisms underlying cortical folding remain debated, sulcal patterns have been linked to differential regional cortical expansion and may therefore indicate relative expansion or reorganisation of cortical territories [43]. Below, we discuss these observations and their potential functional significance, while noting that functional cortical data remain available for only a small number of pinniped species and cortical regions.

**Figure 5.**
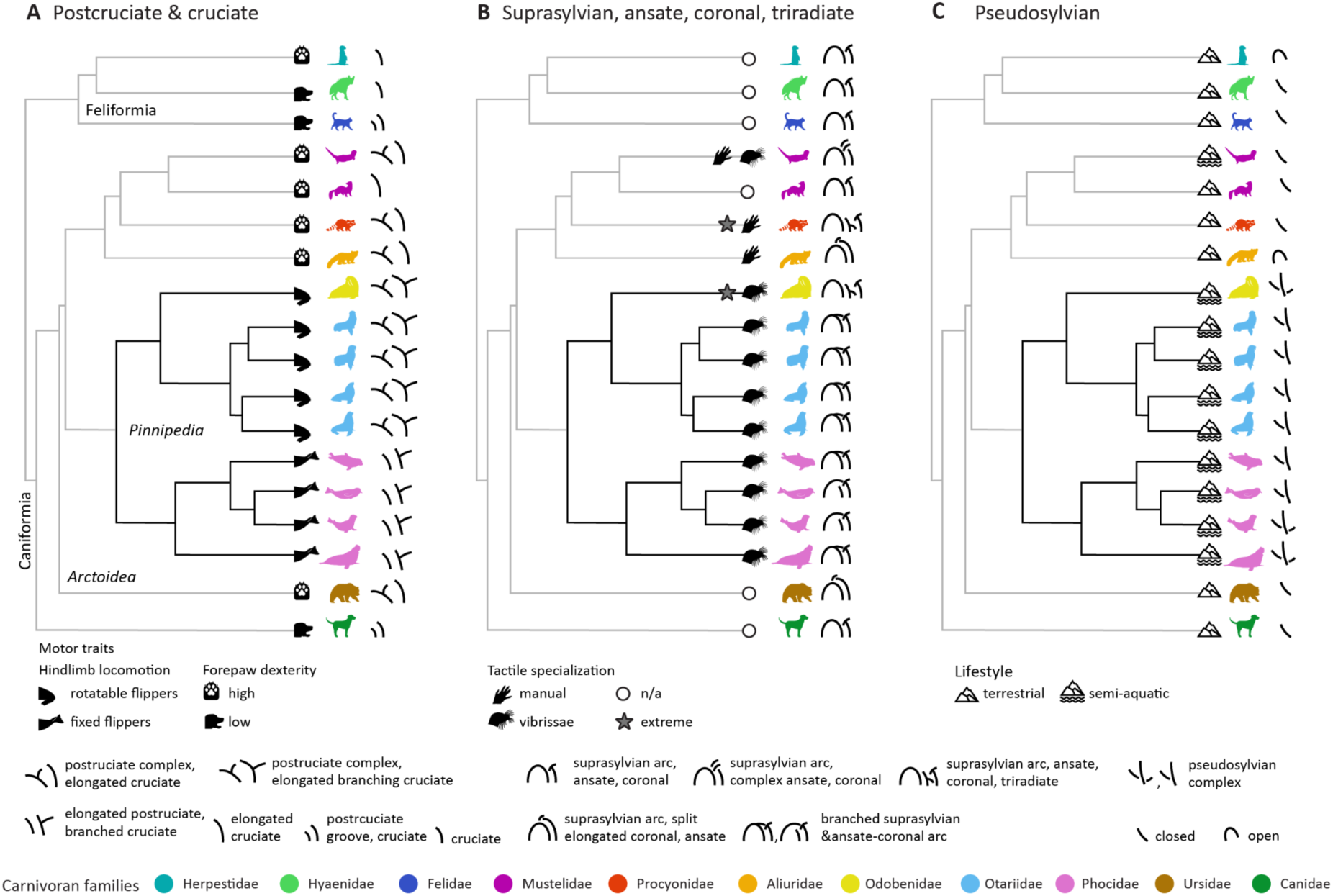
Lineage-specific observations in sulcal organisation and their potential relationship to behaviour and ecology across carnivorans. **(A)** In fissipeds, greater complexity of the postcruciate and cruciate sulci is associated with pronounced forepaw dexterity. Pinnipeds exhibit a different pattern of sulcal variation in this region, potentially related to modifications of somatomotor cortical organisation. Within pinnipeds, these configurations vary alongside differences in hindlimb locomotion between phocids, otariids, and odobenids. (**B)** Sulcal variation in presumed somatosensory territories may relate to forepaw dexterity in fissipeds and reliance on vibrissal sensation in pinnipeds. The semi-aquatic Asian small-clawed otter combines both forms of tactile specialisation. (**C**) Pinnipeds exhibit additional secondary branches surrounding the pseudosylvian fissure, forming a ‘pseudosylvian sulcal complex’. This may reflect modifications of sensory cortical organisation associated with the transition to marine environments. Fissiped sulcal configurations and behavioural observations are based on (Boch et al., 2026); see Figure 6 for information on functional cortical territories. Phylogenetic branches are black for pinnipeds and dark grey for fissipeds.

**Figure 6.**
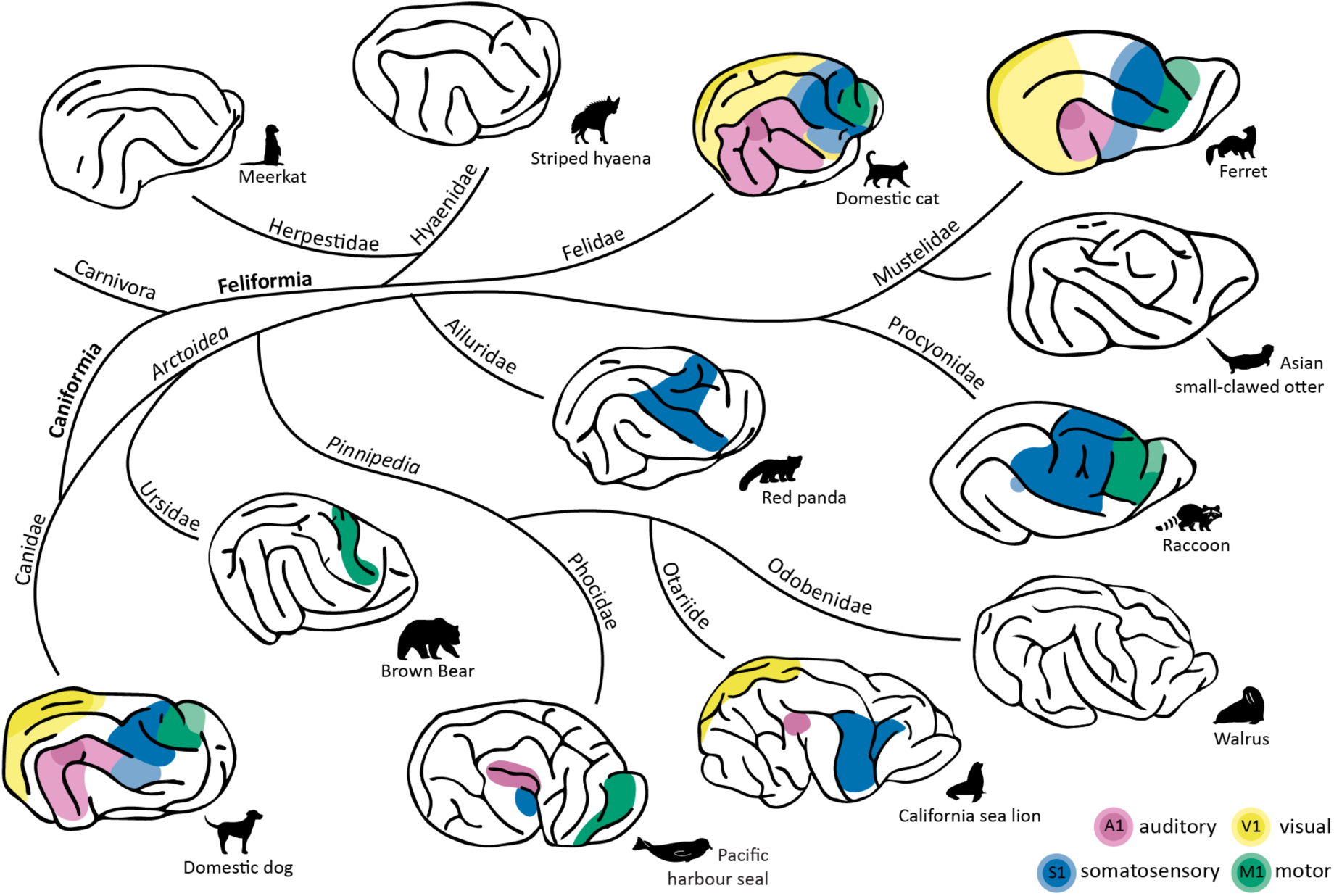
Carnivoran brain diversity illustrated by representatives of eleven families. Schematic lateral views show the approximate locations of primary visual (V1), somatosensory (S1), auditory (A1), and motor (M1) cortical territories in representatives of eleven carnivoran families, combining pinnipeds examined here with fissipeds from [21]. Functional territories are shown where supported by previous anatomical, electrophysiological, or neuroimaging studies; uncoloured territories indicate unavailable or unrepresented functional information; see **Supplementary Figure S4** for corresponding sulcal anatomy and nomenclature.

#### Frontoparietal cortical territory

Variation in sulcal organization across carnivorans was especially apparent in the frontoparietal cortex (**Figures 5A-B**). In fissipeds, increased complexity of the cruciate and postcruciate sulci is associated with pronounced forepaw dexterity, particularly among arctoid species [8,21,40,44]. Pinnipeds exhibited a different pattern of sulcal organisation in this territory: across all three families, the cruciate sulcus exhibited an additional branch compared to fissiped arctoids, while otariids and the walrus additionally showed more elaborate postcruciate configurations. Functional and cytoarchitectonic studies indicate that this broader frontoparietal territory encompasses motor and somatosensory representations across carnivorans [24,26,28,39,44–51,52; see **Figure 6]**.

In the harbour seal, electrical stimulation identified an extensive motor territory rostral and ventral to the cruciate sulcus, with substantial fore- and hind-flipper representations in which stimulation elicited swimming-like movements [24,27; Figure 6]. The transition to aquatic locomotion in pinnipeds involved substantial modifications of the postcranial locomotor system, including the limbs, vertebral column and sacrum [53–55]. The distinct cruciate configuration of pinnipeds relative to fissipeds may therefore reflect corresponding modifications of somatomotor cortical organisation.

Further variation *within* pinnipeds was apparent around the postcruciate and ansate sulci (**Figure 5A**). The available functional studies are consistent with this region containing part of the primary somatosensory cortex (S1). In the northern fur seal, the dorsal portion of S1, in the region of the ansate and postcruciate sulci, contains representations of the hindlimbs and trunk [26]. A broadly corresponding location of S1 has been identified cytoarchitecturally in the California sea lion [28; see Figure 6]. Pinniped families differ substantially in their locomotor strategies: phocids rely predominantly on their hindlimbs for aquatic propulsion and make limited use of them during terrestrial locomotion, whereas otariids use their forelimbs for aquatic propulsion but retain quadrupedal locomotion on land. Walruses combine aspects of both strategies, relying primarily on hindlimb propulsion in water while retaining an otariid-like quadrupedal gait on land [27,53–55]. Thus, whereas phocid locomotion involves comparatively limited limb-based locomotor demands on land, otariids and walruses employ the same limbs in distinct and elaborate locomotor patterns depending on whether they are moving in water or on land. The greater sulcal complexity observed around the postcruciate and ansate sulci in otariid and odobenid species may therefore relate to the somatomotor demands of switching between these different modes of limb use.

Sulcal variation within the frontoparietal territory may also relate to tactile specialisation (**Figure 5B**). In fissipeds, sulcal organisation around the ansate, coronal, and rostral suprasylvian sulci varies with forepaw specialisation, with particularly pronounced complexity in species such as the raccoon, where an expanded forepaw representation is accompanied by an additional triradiate sulcus [39,40,44,56]. Pinnipeds instead rely extensively on specialised vibrissae for tactile exploration and hydrodynamic sensing underwater [57–59]. In the northern fur seal, a substantial representation of the head and vibrissae occupies the territory between the rostral suprasylvian and coronal sulci [26], which is expanded and reoriented in pinnipeds relative to most terrestrial carnivorans, potentially reflecting increased reliance on vibrissal sensation during aquatic foraging [30]. Consistent with this, we found a distinct sulcal configuration in this region: the predominant orientation of the coronal sulcus was approximately orthogonal between pinnipeds and fissipeds (**Figure 6**), and the rostral suprasylvian sulcus was frequently concealed beneath the intervening gyrus, consistent with its greater prominence in pinnipeds. Vibrissal specialisation also varies within pinnipeds, with phocids exhibiting pronounced adaptations for hydrodynamic sensing [60]. The walrus provides a particularly interesting example, as it is highly specialised for whisker-based tactile exploration [57,61] and exhibited an additional, triradiate-shaped sulcus within this presumed vibrissal territory. Although its identification should be treated with caution given the limited specimen quality and availability of only one reconstructed hemisphere, its location raises the possibility of further modification of this somatosensory territory in highly specialised vibrissal species. Together, these observations suggest that this territory may have been modified both with increased reliance on vibrissal sensation in aquatic environments and with further specialisation of vibrissal function within pinnipeds.

A further family-level difference was apparent in the superorbital sulcus, which was present in all phocids examined and in the walrus, but could not be consistently identified in otariids. In the walrus, it additionally exhibited a small secondary branch. Located in the lateral frontal cortex, this sulcus lies within a territory in which Rioch [24] identified motor representations of the head and neck in the harbour seal, and where Cook et al. [62] found direct projections to the nucleus ambiguus, a brainstem nucleus involved in laryngeal motor control. Vocal production learning and broader vocal plasticity have been described in several phocids, including elephant seal [63], grey seal [64], and harbour seal [65], as well as in the walrus, which learns a seasonal mating song [66]. In contrast, otariid vocalisations are highly stereotyped [67]. The more pronounced expression of the superorbital sulcus in phocids and the walrus may therefore reflect greater elaboration of cortical territories associated with enhanced vocal motor control.

#### Temporal cortical territory

Another characteristic feature of pinniped cortical organisation were the additional secondary branches around the pseudosylvian fissure (**Figures 5C** and **6**), contributing to the pseudosylvian complex described above, and contrasting with the comparatively simpler configuration in most fissipeds [21]. Electrophysiological mapping in the harbour seal places auditory cortex in close proximity to the pseudosylvian fissure [25] and recent diffusion MRI analyses similarly locate primary auditory cortex within this territory in the California sea lion and other pinniped species [62; see Figure 6]. Pinnipeds exhibit pronounced auditory adaptations, with phocids and otariids exhibiting broader hearing ranges than most terrestrial mammals [68–70]. This may reflect increased reliance on audition in aquatic environments, particularly under turbid or low-light conditions. Pinnipeds also retain sensitive hearing in air, supported by adaptations of the peripheral auditory system that enable hearing across both aquatic and terrestrial environments [71]. Auditory experience is also central to vocal learning: several pinniped species, including the walrus and several phocids, modify their vocalisations based on exposure to conspecific vocalisations [67], which may relate to increased connectivity between putative auditory and premotor cortical territories in phocids [62]. The pseudosylvian territory may also encompass other sensory specialisations. Alderson et al. [25] identified a separate mystacial somatosensory representation adjacent to the auditory territory in the harbour seal (see **Figure 6**), a configuration that, to our knowledge, has not been described in fissipeds. Together, the increased sulcal complexity surrounding the pseudosylvian fissure may therefore reflect regional modifications of sensory cortex associated with the transition to marine environments.

## 3. General discussion

In the present study, we examined the neocortical sulcal anatomy across nine marine pinniped species representing all three extant families and compared them with our previous observations in fissipeds [21]. This revealed that the major neocortical sulci are broadly conserved across carnivorans, but their configuration varies substantially across lineages (**Figures 5-6**). Within this shared sulcal *Bauplan*, systematic differences in sulcal configuration distinguish pinniped from fissiped carnivorans. Pinniped brains are overall more gyrencephalic than those of fissipeds with comparable brain mass [15] and exhibit in some cortical territories an approximately orthogonal orientation of major sulci.

As in fissipeds, species of the same pinniped family generally exhibited more similar sulcal patterns. Within these family-level configurations, however, we also observed further species-specific modifications. The walrus and raccoon provide notable examples among pinniped and fissiped arctoids, respectively. Both exhibit an additional sulcus in presumed somatosensory territories, potentially related to their respective tactile specialisations [21,56,61,72]. While some of the features described here have been noted previously [e.g., 22,24,28], our broader sample now allows us to integrate these previously isolated observations into a common anatomical framework. Together, these comparisons generate new hypotheses about how cortical organisation has diversified in relation to major behavioural and ecological transitions.

Major ecological transitions provide an opportunity to examine how changes in the sensory and motor demands placed on an animal are reflected in brain organisation. MacIver and Finlay [73] considered this question across the water-to-land transition in vertebrates, highlighting how fundamental changes in the physical environment altered the information available to animals and the sensory and motor problems their nervous systems had to solve. The return of mammals to aquatic environments provides a complementary perspective, with this transition similarly proposed to alter the demands placed on the nervous system [74]. The differences we observed in temporal and presumed somatomotor cortical territories between pinnipeds and fissipeds may reflect changes in sensory and motor demands associated with this transition [75,76].

Otters provide a particularly informative comparison within Carnivora, given their tactile specialisations and distinctive sulcal configurations surrounding the somatosensory cortex [21,77; **Figure 5B**]. Comparative analyses further identify otters as distinct in their brain organisation among musteloids [17]. Sampling species with different degrees of aquatic specialisation could help distinguish cortical features associated with increasing reliance on aquatic environments. Within pinnipeds, increased intraspecific sampling could reveal whether sulcal variation relates to sex-specific differences in movement and foraging ecology. Northern elephant seals, for example, exhibit pronounced sexual dimorphism accompanied by markedly different migration and foraging strategies between males and females [3,78,79]. Integrating such comparisons with complementary neuroanatomical data, including connectivity [62], could further help determine how variation in sulcal anatomy relates to underlying cortical organisation.

Pinnipeds are among the few extant mammalian lineages to have transitioned from terrestrial ancestors into marine environments [80,81]. Cetaceans provide a particularly informative comparison, having transitioned to fully aquatic life from terrestrial artiodactyl ancestors [3,4,80]. Extending the comparative framework established here across independently evolved semi-aquatic and aquatic lineages can help distinguish convergent patterns associated with life in water from lineage-specific modifications. By establishing a common anatomical framework across carnivorans, our findings provide a basis for moving beyond descriptions of brain diversity towards testing how cortical organisation has diversified across major ecological transitions.

## 4. Materials and methods

All data are MRI scans of post-mortem samples, which we harmonised during the initial preprocessing step to account for variations in scanning protocols (**Supplementary Table 1**). For the majority of species, we applied a standardised processing pipeline to reconstruct cortical surfaces of both hemispheres. Two specimens, the walrus and the Guadalupe fur seal, required partial manual surface reconstruction due to image quality issues. Building on the limited previous work and our own observations, we developed detailed descriptions of the major neocortical sulci and formalised criteria for their identification. These criteria build on our previously established framework for fissiped carnivorans [21] and were systematically extended to capture pinniped sulcal anatomy.

### 4.1. Data

The sample comprises nine pinniped species (**Figure 1**). One representative individual per species is presented in the main text, except for the grey seal, where an averaged template is used. To enable assessment of intraspecific variation, additional individuals were included for several species and are reported in the Supplementary material, bringing the total to 16 individuals (**Supplementary Table 1**).

### 4.2. Procedure

All data were obtained from post-mortem brain samples collected from animals that died of natural causes, incidental bycatch, or were euthanised for medical, welfare-related, or management reasons unrelated to this study. All brains were placed in fixative within 24 hours of death for post-mortem imaging. Samples were obtained from multiple research centres; detailed information on sample provenance, fixation and storage procedures, relevant permits, and data sources is provided in the **Extended Materials & Methods** and **Supplementary Table 1**.

### 4.3. Imaging protocols and data preprocessing

MRI data were obtained across multiple research centres using structural or diffusion MRI acquisition protocols. To harmonise data across acquisitions and generate images with T1w-like contrast suitable for cortical surface reconstruction, scans were preprocessed using tools from FSL (https://fsl.fmrib.ox.ac.uk), ANTs [82], and the MR Comparative Anatomy Toolbox (MrCat; https://www.neuroecologylab.org). Detailed acquisition parameters and specimen-specific preprocessing procedures are provided in the **Extended Materials & Methods**.

#### Surface reconstruction

All scans were reoriented to AC-PC alignment and standard FSL orientation. We applied intensity bias-field correction to all structural scans using ANTs. Cortical surfaces were reconstructed primarily using precon_all [83], an adapted version of FreeSurfer’s recon-all pipeline [84] designed for non-human animal models. Due to reduced image quality or damaged tissue, the walrus and Guadalupe fur seal required partial manual reconstruction using the Stitcher pipeline [85]. Irrespective of the reconstruction approach, all final cortical surfaces were converted to GIFTI surface format (.surf.gii). Detailed reconstruction and specimen-specific procedures are provided in the **Extended Materials & Methods**.

#### Protocols for labelling and sulcal masks

We labelled all major neocortical sulci visible on the cortical surface by creating digital sulcal masks on smoothed midthickness surfaces using Connectome Workbench [86]. Labelling was carried out bilaterally, except for the Guadalupe fur seal and walrus, for which only the right hemisphere could be reconstructed. To reduce visual complexity, figures show only right hemispheres and one individual per species; hemispheric and intraspecific variation is addressed in the text and supplementary material.

We initially applied the protocols established for identifying fissiped sulcal anatomy [21, previously referred to as ‘recipes’], drawing particularly on the ursid brain, which appears most similar in overall cortical morphology to pinnipeds among land-dwelling arctoids [21,87,88]. We then began to label pinniped brains for which previous neuroanatomical descriptions were available. We then compared these labels with available descriptions of pinniped brains, which are limited to individual species or cortical regions [22–24,26,28,89,90], a grey seal atlas [38], and comparative analyses of digital cranial endocasts [29,30]. Unless stated otherwise, our descriptions were consistent with previous observations. Based on these previous descriptions and our own observations, we then expanded the fissiped sulcal protocols to accommodate pinniped sulcal anatomy. Anatomical nomenclature primarily follows Czeibert et al. [91]; alternative sulcal names are provided once.

#### Variations in sulcal anatomy and functional considerations

We characterised variation in neocortical sulcal anatomy across pinnipeds and, where relevant, compared these patterns with previously described fissiped brains. Variation was assessed qualitatively based on sulcal presence or absence, continuity, branching patterns, and relative prominence, focusing on interspecific and lineage-level differences; intraspecific variation was examined primarily to confirm the consistency of major sulcal patterns. To facilitate comparisons across carnivoran families, we used schematic drawings from Boch et al. [21] summarising sulcal configurations across eight fissiped families (**Figure 1**), complemented by newly generated drawings for three pinniped representatives.

Where available, we additionally indicated the approximate locations of major sensory and motor cortical regions as a coarse functional reference frame. Information was derived from prior physiological, histological, cytoarchitectural, and diffusion MRI studies, but direct functional mapping of pinniped cortex remains sparse. Functional cortical information is available for only three species in our sample: the Pacific harbour seal, California sea lion, and northern elephant seal. In the harbour seal, electrical stimulation and electrophysiological studies have mapped motor, auditory, and mystacial somatosensory representations [24,25,27], while somatosensory and visual cortical organisation has been described in the California sea lion and visual cortical organisation in the northern elephant seal [28,90]. Diffusion MRI and histological work have further identified cortical and subcortical connectivity associated with vocal motor control and auditory-premotor processing across pinnipeds [62]. Functional interpretations, therefore, remain tentative and are limited to cautious comparisons informed by sulcal topology and available functional information in other carnivorans.

## 5. Data availability statement

All derived data supporting the article, including sulcal masks and associated midthickness cortical surface reconstructions for all 16 animals, are publicly available at https://git.fmrib.ox.ac.uk/neuroecologylab/carnivore-surfaces, along with the same data for 32 fissipeds. The corresponding raw data (T1-like volumetric MRI scans) are available via the Digital Brain Zoo of the University of Oxford [33; https://open.win.ox.ac.uk/DigitalBrainBank/#/datasets/zoo], subject to agreement to a Material Transfer Agreement in accordance with ethical and funding requirements. The grey seal template is openly available at https://osf.io/hz3gc/ [38].

## 6. Acknowledgements

The authors would like to thank veterinarians, pathologists, and technicians at The Marine Mammal Center in Sausalito, CA, who provided California sea lion, harbor seal, and northern elephant seal brain samples from animals who died while in rehabilitation. We also thank Radek Sharma, Louisa Pillwax and Celine Parizek for their assistance in creating the digital sulcus masks.

MB’s and SB’s research was funded in whole or in part by the Austrian Science Fund (FWF) 10.55776/J4828 and 10.55776/PAT4698926, and the Messerli Foundation (Sörenberg, Switzerland). KAS’s research was funded by the Serrapilheira Institute, Grant number: 5057. NP’s research was funded by Japan Society for the Promotion of Science (JSPS), Grant number: 21K06404. PFC’s research was funded in part by a Human Frontiers Science Program Research Grant: https://doi.org/10.52044/HFSP.RGP00192022.pc.gr.153618. The work of RBM is supported by the Medical Research Council (MRC) UK [MR/Y010698/1]. The funders had no role in study design, data collection and analysis, decision to publish, or preparation of the manuscript. For the purpose of Open Access, the authors have applied a CC BY public copyright licence to any Author Accepted Manuscript version arising from this submission.

## References

1. Berta A. 2012 Return to the Sea: The Life and Evolutionary Times of Marine Mammals. University of California Press.

2. Valkenburgh BV. 1999 MAJOR PATTERNS IN THE HISTORY OF CARNIVOROUS MAMMALS. Annual Review of Earth and Planetary Sciences 27, 463–493. (doi:10.1146/annurev.earth.27.1.463)

3. Mittermeier RA, Wilson DE, editors. 2014 Handbook of the mammals of the world. Vol 4: Sea mammals. Barcelona: Lynx Editions.

4. Wilson DE, Mittermeier RA, editors. 2009 Handbook of the mammals of the world. Vol 1: Carnivores. Barcelona: Lynx Editions.

5. Kumar S, Stecher G, Suleski M, Blair Hedges S. 2017 TimeTree: A Resource for Timelines, Timetrees, and Divergence Times. Molecular Biology and Evolution 34, 1812– 1819. (doi:10.1093/MOLBEV/MSX116)

6. Finarelli JA, Flynn JJ. 2009 Brain-size evolution and sociality in Carnivora. Proceedings of the National Academy of Sciences of the United States of America 106, 9345–9349. (doi:10.1073/pnas.0901780106)

7. Holekamp KE, Benson-Amram S. 2017 The evolution of intelligence in mammalian carnivores. Interface Focus 7. (doi:10.1098/RSFS.2016.0108)

8. Iwaniuk AN, Pellis SM, Whishaw IQ. 1999 Brain Size Is Not Correlated with Forelimb Dexterity in Fissiped Carnivores (Carnivora): A Comparative Test of the Principle of Proper Mass. Brain, Behavior and Evolution 54, 167–180. (doi:10.1159/000006621)

9. Kirkwood TMB, Shultz S, Silcox MT, Schwab JA, Palmer SM, Rule JP, Jones KE. 2026 The land-to-water transition impacts brain shape in caniform carnivorans. Evol, qpag024. (doi:10.1093/evolut/qpag024)

10. Michaud M, Toussaint SLD, Gilissen E. 2022 The impact of environmental factors on the evolution of brain size in carnivorans. Communications Biology 5, 1–13. (doi:10.1038/s42003-022-03748-4)

11. Smaers JB, Dechmann DKN, Goswami A, Soligo C, Safi K. 2012 Comparative analyses of evolutionary rates reveal different pathways to encephalization in bats, carnivorans, and primates. Proceedings of the National Academy of Sciences 109, 18006–18011. (doi:10.1073/pnas.1212181109)

12. Smaers JB et al. 2021 The evolution of mammalian brain size. Science Advances 7, eabe2101. (doi:10.1126/SCIADV.ABE2101)

13. Chengetanai S et al. 2020 Brain of the African wild dog. I. Anatomy, architecture, and volumetrics. Journal of Comparative Neurology 528, 3245–3261. (doi:10.1002/CNE.24999)

14. Grewal JS et al. 2020 Brain gyrification in wild and domestic canids: Has domestication changed the gyrification index in domestic dogs? Journal of Comparative Neurology 528, 3209–3228. (doi:10.1002/cne.24972)

15. Manger PR, Prowse M, Haagensen M, Hemingway J. 2012 Quantitative analysis of neocortical gyrencephaly in African elephants (Loxodonta africana) and six species of cetaceans: Comparison with other mammals. Journal of Comparative Neurology 520, 2430–2439. (doi:10.1002/cne.23046)

16. Krubitzer L, Kaas J. 2005 The evolution of the neocortex in mammals: how is phenotypic diversity generated? Current Opinion in Neurobiology 15, 444–453. (doi:10.1016/J.CONB.2005.07.003)

17. Michaud M, Harrington AR, Ladeveze S, Gálvez-López E, Fischer V, Alfsen A, Bertrand O. 2026 Brain scaling and diversity in musteloids. Journal of Anatomy (doi:10.1111/joa.70226)

18. Amiez C, Sallet J, Hopkins WD, Meguerditchian A, Hadj-Bouziane F, Ben Hamed S, Wilson CRE, Procyk E, Petrides M. 2019 Sulcal organization in the medial frontal cortex provides insights into primate brain evolution. Nat Commun 10, 3437. (doi:10.1038/s41467-019-11347-x)

19. Amiez C et al. 2023 A revised perspective on the evolution of the lateral frontal cortex in primates. Science Advances 9. (doi:10.1126/sciadv.adf9445)

20. Connolly C J. 1950 External morphology of the primate brain. Springfield, Ill: Thomas.

21. Boch M et al. 2026 Between-species variation in neocortical sulcal anatomy of the carnivoran brain. eLife 13:RP100851. (10.7554/eLife.100851.3)

22. Fish PA. 1898 The brain of the fur seal, callorhinus ursinus; with a comparative description of those of zalophus californianus, phoca vitulina, ursus americanus and monachus tropicalis. Journal of Comparative Neurology 8, 57–91. (doi:10.1002/cne.910080105)

23. Montie EW, Pussini N, Schneider GE, Battey TWK, Dennison S, Barakos J, Gulland F. 2009 Neuroanatomy and Volumes of Brain Structures of a Live California Sea Lion (Zalophus californianus) From Magnetic Resonance Images. The Anatomical Record 292, 1523–1547. (doi:10.1002/ar.20937)

24. Rioch DMcK. 1937 A Physiological and Histological Study of the Frontal Cortex of the Seal (Phoca vitulina). Biological Bulletin 73, 591–602. (doi:10.2307/1537617)

25. Alderson AM, Diamantopoulos E, Downman CBB. 1960 Auditory cortex of the seal (Phoca vitulina). J Anat 94, 506–511.

26. Ladygina TF, Popov VV, Supin AYa. 1985 Topical organization of somatic projections in the fur seal cerebral cortex. Neurophysiology 17, 246–252. (doi:10.1007/BF01052461)

27. Langworthy OR, Hesser FH, Kolb LC, Rathbone HB, Rathbone JL. 1938 A physiological study of the cerebral cortex of the hair seal (Phoca vitulina). Journal of Comparative Neurology 69, 351–369. (doi:10.1002/cne.900690302)

28. Sawyer EK, Turner EC, Kaas JH. 2016 Somatosensory brainstem, thalamus, and cortex of the California sea lion (Zalophus californianus). Journal of Comparative Neurology 524, 1957–1975. (doi:10.1002/cne.23984)

29. Loza CM, Sánchez-Villagra MR, Scarano AC, Romero M, Barbeito CG, Carlini AA. 2023 The brain of fur seals, seals, and walrus (Pinnipedia): A comparative anatomical and phylogenetic study of cranial endocasts of semiaquatic mammals. J Mammal Evol 30, 1011–1028. (doi:10.1007/s10914-023-09679-z)

30. Lyras GA, Werdelin L, van der Geer BGM, van der Geer AAE. 2023 Fossil brains provide evidence of underwater feeding in early seals. Communications Biology 2023 6:1 6, 1–8. (doi:10.1038/s42003-023-05135-z)

31. Friedrich P et al. 2021 Imaging evolution of the primate brain: the next frontier? NeuroImage 228, 117685. (doi:10.1016/J.NEUROIMAGE.2020.117685)

32. Mars RB, Neubert FX, Verhagen L, Sallet J, Miller KL, Dunbar RIM, Barton RA. 2014 Primate comparative neuroscience using magnetic resonance imaging: promises and challenges. Frontiers in Neuroscience 8, 298. (doi:10.3389/fnins.2014.00298)

33. Tendler BC et al. 2022 The Digital Brain Bank, an open access platform for post-mortem datasets. eLife 11, e73153. (doi:10.7554/eLife.73153)

34. Thiebaut de Schotten M, Croxson PL, Mars RB. 2019 Large-scale comparative neuroimaging: Where are we and what do we need? Cortex 118, 188–202. (doi:10.1016/J.CORTEX.2018.11.028)

35. Doronina L, Churakov G, Shi J, Brosius J, Baertsch R, Clawson H, Schmitz J. 2015 Exploring Massive Incomplete Lineage Sorting in Arctoids (Laurasiatheria, Carnivora). Molecular Biology and Evolution 32, 3194–4204. (doi:10.1093/molbev/msv188)

36. Field MA et al. 2022 The Australian dingo is an early offshoot of modern breed dogs. Science Advances 8, 5944. (doi:10.1126/sciadv.abm5944)

37. Freedman AH et al. 2014 Genome Sequencing Highlights the Dynamic Early History of Dogs. PLOS Genetics 10, e1004016. (doi:10.1371/JOURNAL.PGEN.1004016)

38. Hoeksema N et al. 2021 Neuroanatomy of the grey seal brain: bringing pinnipeds into the neurobiological study of vocal learning. Philosophical Transactions of the Royal Society B: Biological Sciences 376, 20200252. (doi:10.1098/rstb.2020.0252)

39. Welker WI, Seidenstein S. 1959 Somatic sensory representation in the cerebral cortex of the racoon (Procyon lotor). Journal of Comparative Neurology 111, 469–501. (doi:10.1002/CNE.901110306)

40. Welker WI, Campos GB. 1963 Physiological significance of sulci in somatic sensory cerebral cortex in mammals of the family procyonidae. Journal of Comparative Neurology 120, 19–36. (doi:10.1002/CNE.901200103)

41. Kawamura K. 1971 Variations of the cerebral sulci in the cat. Acta Anatomica 80, 204–221. (doi:10.1159/000143689)

42. Radinsky LB. 1969 Outlines of canid and felid brain evolution. Annals of the New York Academy of Sciences 167, 277–288. (doi:10.1111/J.1749-6632.1969.TB20450.X)

43. Ronan L, Fletcher PC. 2015 From genes to folds: a review of cortical gyrification theory. Brain Struct Funct 220, 2475–2483. (doi:10.1007/s00429-014-0961-z)

44. Welker WI, Johnson JI, Pubols BH. 1964 Some Morphological and Physiological Characteristics of the Somatic Sensory System in Raccoons. American Zoologist 4, 75– 94.

45. Chengetanai S, Bhagwandin A, Bertelsen MF, Hård T, Hof PR, Spocter MA, Manger PR. 2025 The Brain of the African Wild Dog. V. The Somatosensory System and Vestibular Nuclear Complex. Journal of Comparative Neurology 533, e70071. (doi:10.1002/cne.70071)

46. Chengetanai S, Bhagwandin A, Bertelsen MF, Hård T, Hof PR, Spocter MA, Manger PR. 2025 The Brain of the African Wild Dog. VI. The Motor System. Journal of Comparative Neurology 533, e70072. (doi:10.1002/cne.70072)

47. Dykes RW, Rasmusson DD, Hoeltzell PB. 1980 Organization of primary somatosensory cortex in the cat. Journal of Neurophysiology 43, 1527–1546. (doi:10.1152/jn.1980.43.6.1527)

48. Gorska T. 1974 Functional organization of cortical motor areas in adult dogs and puppies. Acta Neurobiologiae Experimentalis 34, 171–203.

49. Guran C-NA, Boch M, Sladky R, Lonardo L, Karl S, Huber L, Lamm C. 2024 Functional mapping of the somatosensory cortex using noninvasive fMRI and touch in awake dogs. Brain Struct Funct (doi:10.1007/s00429-024-02798-0)

50. Hardin WB, Arumugasamy N, Douglas Jameson H. 1968 Pattern of localization in ‘precentral’ motor cortex of raccoon. Brain Research 11, 611–627. (doi:10.1016/0006-8993(68)90149-2)

51. Smith WK. 1933 Motor cortex of the bear (Ursus americanus): a physiologic and histologic study. Arch NeurPsych 30, 14–39. (doi:10.1001/archneurpsyc.1933.02240130022002)

52. Tanaka D, Gorska T, Dutkiewicz K. 1981 Corticostriate projections from the primary motor cortex in the dog. Brain Research 209, 287–303. (doi:10.1016/0006-8993(81)90154-2)

53. Esteban JM, Martín-Serra A, Pérez-Ramos A, Rybczynski N, Pastor FJ, Figueirido B. 2023 Investigating the land-to-sea transition in carnivorans from the evolution of sacrum morphology in pinnipeds. J Mammal Evol 30, 341–362. (doi:10.1007/s10914-023-09650-y)

54. Kuhn C, Frey E. 2012 Walking like caterpillars, flying like bats—pinniped locomotion. Palaeobio Palaeoenv 92, 197–210. (doi:10.1007/s12549-012-0077-5)

55. Pierce SE, Clack JA, Hutchinson JR. 2011 Comparative axial morphology in pinnipeds and its correlation with aquatic locomotory behaviour. Journal of Anatomy 219, 502–514. (doi:10.1111/j.1469-7580.2011.01406.x)

56. Iwaniuk AN, Whishaw IQ. 1999 How skilled are the skilled limb movements of the raccoon (*Procyon lotor*)? Behavioural Brain Research 99, 35–44. (doi:10.1016/S0166-4328(98)00067-9)

57. Dougill G, Brassey CA, Starostin EL, Andrews H, Kitchener A, van der Heijden GHM, Goss VGA, Grant RA. 2023 Describing whisker morphology of the Carnivora. Journal of Morphology 284, e21628. (doi:10.1002/jmor.21628)

58. Grant RA. 2026 Vibrissal sensing in mammals in a changing world. J Exp Biol 229, jeb250776. (doi:10.1242/jeb.250776)

59. Zheng X, Kamat AM, Cao M, Kottapalli AGP. 2021 Creating underwater vision through wavy whiskers: a review of the flow-sensing mechanisms and biomimetic potential of seal whiskers. J. R. Soc. Interface. 18, 20210629. (doi:10.1098/rsif.2021.0629)

60. Hanke W, Witte M, Miersch L, Brede M, Oeffner J, Michael M, Hanke F, Leder A, Dehnhardt G. 2010 Harbor seal vibrissa morphology suppresses vortex-induced vibrations. J Exp Biol 213, 2665–2672. (doi:10.1242/jeb.043216)

61. Milne AO, Smith C, Orton LD, Sullivan MS, Grant RA. 2020 Pinnipeds orient and control their whiskers: a study on Pacific walrus, California sea lion and Harbor seal. J Comp Physiol A 206, 441–451. (doi:10.1007/s00359-020-01408-8)

62. Cook PF, Rouse AA, Sawyer E, Miller K, Berns G. 2026 Seal and sea lion brains have evolved to support volitional control of vocal behavior and learning. Science 391, 1146– 1150. (doi:10.1126/science.adx9367)

63. Sanvito S, Galimberti F, Miller EH. 2007 Observational Evidences of Vocal Learning in Southern Elephant Seals: a Longitudinal Study. Ethology 113, 137–146. (doi:10.1111/j.1439-0310.2006.01306.x)

64. Stansbury AL, Janik VM. 2019 Formant Modification through Vocal Production Learning in Gray Seals. Current Biology 29, 2244–2249.e4. (doi:10.1016/j.cub.2019.05.071)

65. Goncharova M, Jadoul Y, Reichmuth C, Fitch WT, Ravignani A. 2024 Vocal tract dynamics shape the formant structure of conditioned vocalizations in a harbor seal. Annals of the New York Academy of Sciences 1538, 107–116. (doi:10.1111/nyas.15189)

66. Sjare B, Stirling I, Spencer C. 2003 Structural variation in the songs of Atlantic walruses breeding in the Canadian High Arctic. Aquatic Mammals 29, 297–319.

67. Reichmuth C, Casey C. 2014 Vocal learning in seals, sea lions, and walruses. Current Opinion in Neurobiology 28, 66–71. (doi:10.1016/j.conb.2014.06.011)

68. Cunningham KA, Reichmuth C. 2016 High-frequency hearing in seals and sea lions. Hearing Research 331, 83–91. (doi:10.1016/j.heares.2015.10.002)

69. Kastak D, Schusterman RJ. 1998 Low-frequency amphibious hearing in pinnipeds: Methods, measurements, noise, and ecology. J. Acoust. Soc. Am. 103, 2216–2228. (doi:10.1121/1.421367)

70. Reichmuth C, Holt MM, Mulsow J, Sills JM, Southall BL. 2013 Comparative assessment of amphibious hearing in pinnipeds. J Comp Physiol A 199, 491–507. (doi:10.1007/s00359-013-0813-y)

71. Wartzok D, Ketten DR. 1999 Marine mammal sensory systems. In Biology of marine mammals (eds JR Thomas, RA Kastelein, AYa Supin), pp. 117–175.

72. Welker WI, Seidenstein S. 1959 Somatic sensory representation in the cerebral cortex of the racoon (Procyon lotor). Journal of Comparative Neurology 111, 469–501. (doi:10.1002/CNE.901110306)

73. MacIver MA, Finlay BL. 2021 The neuroecology of the water-to-land transition and the evolution of the vertebrate brain. Phil. Trans. R. Soc. B 377, 20200523. (doi:10.1098/rstb.2020.0523)

74. Bauer GB, Cook PF, Harley HE. 2020 The Relevance of Ecological Transitions to Intelligence in Marine Mammals. Front. Psychol. 11. (doi:10.3389/fpsyg.2020.02053)

75. Hanke FD, Reichmuth C, Cook P. 2021 The Sensory World of Otariids. In Ethology and Behavioral Ecology of Otariids and the Odobenid (eds C Campagna, R Harcourt), pp. 305–321. Cham: Springer International Publishing. (doi:10.1007/978-3-030-59184-7_16)

76. Hanke FD, Reichmuth C. 2022 Phocid Sensory Systems and Cognition. In Ethology and Behavioral Ecology of Phocids (eds DP Costa, EA McHuron), pp. 31–68. Cham: Springer International Publishing. (doi:10.1007/978-3-030-88923-4_2)

77. Radinsky LB. 1968 Evolution of somatic sensory specialization in otter brains. Journal of Comparative Neurology 134, 495–505. (doi:10.1002/CNE.901340408)

78. Kienle SS, Friedlaender AS, Crocker DE, Mehta RS, Costa DP. 2022 Trade-offs between foraging reward and mortality risk drive sex-specific foraging strategies in sexually dimorphic northern elephant seals. R. Soc. Open Sci. 9, 210522. (doi:10.1098/rsos.210522)

79. Le Boeuf BJ, Crocker DE, Costa DP, Blackwell SB, Webb PM, Houser DS. 2000 Foraging Ecology of Northern Elephant Seals. Ecological Monographs 70, 353–382. (doi:10.1890/0012-9615(2000)070[0353:FEONES]2.0.CO;2)

80. Berta A, Sumach JL, Kovacs KM. 2007 Marine mammals: evolutionary biology (second edition). Polar Research 26, 86–87. (doi:10.3402/polar.v26i1.6210)

81. Uhen MD. 2007 Evolution of marine mammals: Back to the sea after 300 million years. The Anatomical Record 290, 514–522. (doi:10.1002/ar.20545)

82. Avants BB, Tustison N, Johnson H. 2009 Advanced Normalization Tools (ANTS). Insight J 2, 1–35.

83. Benn RA et al. 2025 Precon_all: A species-agnostic automated pipeline for non-human cortical surface reconstruction., 2025.04.16.649072. (doi:10.1101/2025.04.16.649072)

84. Fischl B. 2012 FreeSurfer. NeuroImage 62, 774–781. (doi:10.1016/j.neuroimage.2012.01.021)

85. Mynssen H, Avelino-de-Souza K, Chaim K, Ribeiro VL, Patzke N, Mota B. 2024 Stitcher: A Surface Reconstruction Tool for Highly Gyrified Brains. Neuroinform 22, 539–554. (doi:10.1007/s12021-024-09678-2)

86. Marcus D et al. 2011 Informatics and Data Mining Tools and Strategies for the Human Connectome Project. Frontiers in Neuroinformatics 5.

87. Sienkiewicz T, Sergiel A, Huber D, Maślak R, Wrzosek M, Podgórski P, Reljić S, Paśko Ł. 2019 The Brain Anatomy of the Brown Bear (Carnivora, Ursus arctos L., 1758) Compared to That of Other Carnivorans: A Cross-Sectional Study Using MRI. Frontiers in Neuroanatomy 13, 79. (doi:10.3389/fnana.2019.00079)

88. Smith WK. 1933 Cerebral hemispheres of the american black bear (Ursus americanus): morphologic and phylogenetic characteristics. Archives of Neurology & Psychiatry 30, 1–13. (doi:10.1001/ARCHNEURPSYC.1933.02240130009001)

89. Graïc J-M, Mazzariol S, Casalone C, Petrella A, Gili C, Gerussi T, Orekhova K, Centelleghe C, Cozzi B. 2023 Report on the brain of the monk seal (Monachus monachus, Hermann, 1779). Anatomia, Histologia, Embryologia 53, e12986. (doi:10.1111/ahe.12986)

90. Turner EC, Sawyer EK, Kaas JH. 2017 Optic nerve, superior colliculus, visual thalamus, and primary visual cortex of the northern elephant seal (Mirounga angustirostris) and California sea lion (Zalophus californianus). Journal of Comparative Neurology 525, 2109– 2132. (doi:10.1002/cne.24188)

91. Czeibert K, Piotti P, Petneházy Ö, Kubinyi E. 2018 Sulci of the canine brain: a review of terminology. bioRxiv, 374744. (doi:10.1101/374744)

