## Supplementary Material for "A comparative framework for neocortical sulcal anatomy in pinnipeds"

### Supplementary Material
### A comparative framework for neocortical sulcal anatomy in pinnipeds

Magdalena Boch[1,2,3]*, Kamilla Avelino-de-Souza[4,5]*, Sara Binder[1,2], Nina Patzke[6,7], R. Austin Benn[3], Heitor Mynssen[4,5], Khallil Taverna Chaim[8], Khin Khin Tha[9], Bridget Wicinski[10], Cheuk Y. Tang[11], Paul R. Manger[12], Andrew A. Rouse[13,14], Patrick R. Hof[10], Peter F. Cook[13,14], Rogier B. Mars[3,15]

[1] Department of Cognition, Emotion, and Methods in Psychology, Faculty of Psychology, University of Vienna, Vienna, Austria
[2] Messerli Research Institute, University of Veterinary Medicine Vienna, Medical University of Vienna and University of Vienna, Vienna, Austria
[3] Oxford Centre for Integrative Neuroimaging, FMRIB Centre, Nuffield Department of Clinical Neurosciences, University of Oxford, Oxford, United Kingdom
[4] Brazilian Neurobiodiversity Network, Physics Institute, Federal University of Rio de Janeiro, Rio de Janeiro, Brazil
[5] Department of Biomedical Sciences, Federal University of Rio de Janeiro, Rio de Janeiro, Rio de Janeiro, Brazil
[6] Department of Biological Science, Faculty of Sciences, Hokkaido University, Kita-10, Nishi-8, Sapporo, 060-0810, Japan
[7] Faculty of Medicine, IMBB, HMU Health and Medical University, Olympischer Weg 1, 14471, Potsdam, Germany
[8] LIM44, Hospital das Clinicas HCFMUSP, Faculdade de Medicina, Universidade de Sao Paulo, Sao Paulo, SP, BR
[9] Global Center for Biomedical Science and Engineering, Faculty of Medicine, Hokkaido University, Sapporo, Hokkaido, Japan
[10] Nash Family Department of Neuroscience and Friedman Brain Institute, Icahn School of Medicine at Mount Sinai, New York, New York, USA
[11] Department of Radiology and BioMedical Engineering and Imaging Institute, Icahn School of Medicine at Mount Sinai, New York, New York, USA
[12] Department of Anatomical Sciences, School of Biomedical Sciences, Faculty of Health Sciences, University of the Witwatersrand, Johannesburg, South Africa
[13] Marine Mammal Science and Neuroscience, New College of Florida, United States
[14] Institute for Marine Sciences, University of California Santa Cruz, Santa Cruz, CA, USA
[15] Donders Institute for Brain, Cognition and Behaviour, Radboud University Nijmegen, Nijmegen, Netherlands

* shared authorship and corresponding authors:

Magdalena Boch:
Kamilla Avelino-de-Souza:

#### 1. Extended Materials & Methods

##### 1.1. Procedure

All brains were processed within 24 hours of death and obtained from deceased animals or animals euthanised for medical or welfare-related reasons unrelated to this study. The samples were obtained from multiple research centres (see **Supplementary table 1** for data sources).

California sea lion, harbor seal, and elephant seal brain samples from the New College of Florida brain collection were obtained opportunistically from animals who died of natural causes or were euthanized in marine mammal rehabilitation centers. Brains were cold-fixed in 10% buffered formalin solution. After a minimum of four weeks of fixation, samples were moved to gadolinium-doped agar for scanning (as detailed in 35). The Guadalupe fur seal brain underwent the same fixation procedure but was scanned in a fomblin solution (36).

Steller sea lion brain samples from Hokkaido University were collected at the Shakotan Peninsula, Hokkaido, Japan, and euthanised as part of pest control measures. Spotted seal brains were collected post-mortem following incidental bycatch in fishing nets. The brains were placed in 4% paraformaldehyde (PFA) within 12 hours of death and then transferred to a solution of 0.1 M phosphate-buffered saline (PBS) with 0.1% sodium azide solution and stored at 4°C before scanning.

The Subantarctic fur seal brain from the Brazilian Neurobiodiversity Network was collected at the Instituto ORCA facilities in Guarapari, Espírito Santo, Brazil. The collection was conducted under the Biodiversity Information and Authorization System (SISBIO) permit No 64724-13 and the authorization for collection and transport of biological materials (ABIO) permit No 861/2017, respectively. The brain was removed immediately after death and fixed in 4% PFA for 3 days, after which it was transferred to a 0.1 M PBS solution containing 0.1% sodium azide and stored at 4 °C before scanning.

The grey seal brain template (37) was generated from two post-mortem grey seal brains obtained from animals that died of natural causes or were euthanised for welfare reasons unrelated to the study. Brains were extracted within 2 hours post-mortem, fixed in 10% buffered formalin for approximately six weeks, suspended in 2% agar before scanning to reduce image artefacts, and scanned under stable room temperature conditions.

The walrus brain (Icahn School of Medicine at Mount Sinai collection) was obtained from a specimen from the New York Aquarium (Coney Island) and extracted within 9 hours post-mortem at the Bronx Zoo pathology department, during which the bilateral frontal poles sustained damage. The brain was immersion-fixed in formaldehyde (10% buffered formalin) and subsequently stored in 4% PFA until post-mortem imaging.

**Supplementary Table 1**

Overview Carnivoran Brain Collection - Pinnipeds

| Common name | family | genus | species | sex | age | scan | data |
|---|---|---|---|---|---|---|---|
| Walrus[a] | Odobenidae | *Odobenus* | *rosmarus* | m | A | 3 T | 6 |
| California sea lion 1 | Otariidae | *Zalophus* | *californianus* | m | J | 3 T | 1 |
| California sea lion 2 | Otariidae | *Zalophus* | *californianus* | f | A | 3 T | 1 |
| California sea lion 3 | Otariidae | *Zalophus* | *californianus* | m | J | 3 T | 1 |
| Guadalupe fur seal[a] | Otariidae | *Arctocephalus* | *townsendi* | m | A | 1.5 T | 5 |
| Steller sea lion 1 | Otariidae | *Eumetopias* | *jubatus* | m | J | 3 T | 3 |
| Steller sea lion 2 | Otariidae | *Eumetopias* | *jubatus* | m | A | 3 T | 3 |
| Steller sea lion 3 | Otariidae | *Eumetopias* | *jubatus* | m | A | 3 T | 3 |
| Subantarctic fur seal | Otariidae | *Arctocephalus* | *tropicalis* | m | J | 7 T | 2 |
| Grey seal | Phocidae | *Halichoerus* | *grypus* | n/a | n/a | 3 T | 4 |
| Northern elephant seal 1 | Phocidae | *Mirounga* | *angustirostris* | m | J | 3 T | 1 |
| Northern elephant seal 2 | Phocidae | *Mirounga* | *angustirostris* | m | J | 3 T | 1 |
| Pacific harbour seal 1 | Phocidae | *Phoca* | *vitulina* | f | J | 3 T | 1 |
| Pacific harbour seal 2 | Phocidae | *Phoca* | *vitulina* | m | J | 3 T | 1 |
| Spotted seal 1 | Phocidae | *Phoca* | *largha* | m | A | 3 T | 3 |
| Spotted seal 2 | Phocidae | *Phoca* | *largha* | m | A | 3 T | 3 |

*Note.* The sample comprises 16 individuals from 9 species. The animals belong to the carnivoran suborder Caniformia and include representatives of all three pinniped families: Odobenidae, Otariidae (eared seals and sea lions), and Phocidae (true seals). All data are post-mortem samples obtained from multiple sources: 1 = New College of Florida (Florida, USA), 2 = Brazilian Neurobiodiversity Network, 3 = Hokkaido University (Hokkaido, Japan), 4 = volumetric template (37), 5 = SeaWorld of California (SWC) Marine Mammal Rescue Center for supportive care and rehabilitation, 6 = Icahn School of Medicine at Mount Sinai (New York, USA). The field strength of the scanners was 1.5, 3, or 7 Tesla (T; see scan protocol). For consistency, only the first individual per species is described in the main text; additional individuals are included for comparison in the supplementary material. [a] We only reconstructed the right hemisphere due to physical damage during brain extraction (walrus) or due to algal excitotoxic damage (Guadalupe fur seal) of the left. A, adult; J, juvenile; f, female; m, male; n/a, not applicable.

#### 1.2. Imaging protocols and data preprocessing

The majority of data were obtained using 3 T human MRI scanners, except for the Guadalupe fur seal (1.5 T) and the subantarctic fur seal (7 T). The dataset includes structural and diffusion MRI protocols. To homogenise data across acquisition protocols and generate images with a T1w-like contrast suitable for surface reconstruction, all scans were preprocessed using tools from FSL (https://fsl.fmrib.ox.ac.uk), ANTs (42), and the MR Comparative Anatomy Toolbox (MrCat; https://www.neuroecologylab.org).

All New College of Florida Brain Collection samples were scanned on a 3 T Siemens Trio MRI system (Siemens HealthCare, Erlangen, Germany) using a 32-channel human head receive coil. High-resolution structural images were acquired using a TRUFI (true fast imaging with steady-state free precession) sequence, and diffusion MRI data were collected for all specimens (see Cook et al., 2026 for detailed acquisition parameters and diffusion MRI preprocessing). Diffusion MRI data were acquired using a steady-state free precession (SSFP) sequence (Miller et al., 2012) with 1 mm isotropic resolution, a Field of View (FoV) of 128 mm; TR (repetition time) /TE (echo time) = 31/24 ms; flip angle = 35°; bandwidth = 159 Hz/pixel; q = 10 $cm^{-1}$ and standard gradients. Structural TRUFI scans were acquired at a spatial resolution of 0.6 × 0.6 × 0.5 $mm^3$ (TR/TE = 7.09/3.55 ms; flip angle = 37°). T1-like contrast images required for cortical surface reconstruction were generated either from diffusion MRI data (California sea lions 2-3, northern elephant seals) or from the structural TRUFI scans (Pacific harbour seals, California sea lion 1).

Magnetic resonance imaging of extracted brains from three Steller sea lions (SSL1-SSL3) and two spotted seals (SS1-SS2) was performed on a 3 T MRI system (Magnetom Prisma, Siemens HealthCare, Erlangen, Germany) using a 64-channel head coil. Except for SSL3, all specimens were imaged with a T2-Sampling Perfection with Application-optimized contrast using different flip angle Evolution (T2-SPACE) sequence using the following parameters: axial acquisition; repetition time/echo time (TR/TE) = 4000/410 ms; refocusing angle (RFA)= 120; isotropic spatial resolution of 0.78 mm (field of view (FOV) = 237 × 200 $mm^2$; matrix size = 304 × 256); echo train length = 246; bandwidth= 700 Hz/pixel; number of signal averages = 1; the number of slices = 320, and acquisition time = 12 minutes 24 seconds. SSL3 was imaged using magnetization prepared two rapid acquisition of gradient echoes (MP2RAGE) sequence using the following parameters: sagittal acquisition; TR/TE = 5000/3.08 ms; inversion times (TI1 and TI2) = 935 ms and 2770 ms; flip angle = 5; isotropic spatial resolution of 0.86 mm (FOV = 261 × 220 $mm^2$; matrix size = 304 × 256); echo train length = 1; bandwidth = 240 Hz/pixel; number of signal averages = 1; the number of slices = 224, and acquisition time = 13 minutes 51 seconds.

Guadalupe fur seal brain data were obtained on a Siemens Tim/TRIO 3 T MRI system (Siemens HealthCare, Erlangen, Germany) using a 32-channel receive-only coil. The T1-weighted structural scan was acquired using an MPRAGE sequence with .8 mm isotropic spatial resolution (TR/TE = 2.3/3.3 ms; inversion time = .7 s; flip angle = 6°. We were only able to reconstruct the right hemisphere due to algal excitotoxic damage on the left hemisphere (see Schmitt et al., 2023 for more detailed information).

MRI structural scans of the Subantarctic fur seal were acquired in a 7 T Siemens Classic Magneton scanner (Siemens Healthcare, Erlangen, Germany) equipped with a 32-channel head coil with single radiofrequency transmission (Nova Medical, Wilmington, USA) at the "Image Platform in the Autopsy Room" (PISA) facilities of the School of Medicine at the University of São Paulo, in Brazil. A 3D Gradient Echo Fast Low Angle Shot sequence (3D-FLASH) was acquired with the following scan parameters: TR/TE = 28/4 ms; .5 mm isotropic spatial resolution (FoV = 128 × 128 mm; matrix 256× 256, 128 slices), flip angle = 10°; NSA = 1; BW = 501 Hz/px. The acquisition time was 15 minutes and 21 seconds.

For the grey seal, we used a public template (37) generated from post-mortem MRI scans of two individuals. The scans were acquired on a 3T Philips Intera MRI scanner (Philips Medical Systems, Best, the Netherlands) equipped with a 32-channel head coil. T2-weighted fluid inversion recovery (T2 FLAIR) images were acquired with the following parameters: TR/TE = 11000/107 ms; spatial resolution of 0.86 × 0.86 × 2 $mm^3$; field of view (FOV) = 220 × 220 $mm^2$; matrix size = 256 × 256; flip angle = 90°; number of slices = 69; acquisition time = 13 minutes 45 seconds. The individual transverse T2-FLAIR images were used to generate the anatomical template, which had a final isotropic voxel resolution of 0.86 mm.

MRI data for the walrus brain were obtained using a 3 T Siemens Allegra MRI system (Siemens Healthcare, Erlangen, Germany) at the Icahn School of Medicine at Mount Sinai. The high-resolution T2-weighted anatomical volume of the whole brain was acquired with a turbo spin echo pulse sequence. The imaging protocol was as follows: TR/TE = 6.3/ 75 ms, number of signal averages 10, slice thickness 2 mm, gap 0, matrix size 256 x 256, and FOV 17.9 cm.

##### 1.3. Surface reconstruction

Voxel intensities of the TRUFI and T2-weighted structural scans were inverted (multiplied by -1), and cerebrospinal fluid (CSF) intensities were set to zero to generate T1-like contrast images required for cortical surface reconstruction. For diffusion MRI data, T1-like contrast images were generated by summing the mean_f{1,2,3}_samples images representing estimated fibre fractions across fibre populations at each voxel, applying a square-root intensity transform, scaling by a factor of 2, and masking with the diffusion brain mask.

We then created individual brain masks for all scans to remove remaining non-brain tissue using ITK-SNAP's segmentation tool (44) and manually refined them where necessary.

Cortical surface meshes were generated primarily using precon_all (40; https://github.com/neurabenn/precon_all), an adapted version of FreeSurfer's recon-all pipeline (41) designed for non-human animal models. Due to reduced image quality (e.g., weak contrast or partially damaged tissue) the walrus and Guadalupe fur seal surfaces were manually delineated and reconstructed using the Stitcher pipeline (42) or a combination of Stitcher and precon_all.

For precon_all, we created hemisphere-specific masks using FSLeyes and additional "subcortical" and "noncortical" masks using ITK-SNAP (see 40, for a detailed description of the pipeline). The cerebellum, brainstem, and olfactory bulb were included in the noncortical mask and excluded from the cortical surfaces. Where necessary, we manually refined white matter masks after the initial precon_all run to further improve surface reconstruction. Precon_all was then rerun to generate white, pial, and mid-thickness surfaces, which were subsequently downsampled to 10,000 vertices. Finally, surfaces were spatially smoothed using Connectome Workbench tools (43) with a smoothing strength of 0.2 (15 iterations) to facilitate sulcal labelling.

For Stitcher, we manually traced contours corresponding to the pial surface and, when visible, the white-matter boundary using the OsiriX tracing tool (45). Then, the Stitcher Python library transforms the stack of 2D contours into a closed 3D surface. For the Guadalupe fur seal, because there was no white-matter boundary contrast, we smoothed the pial surface with a strength of 1 and 100 total iterations using Connectome Workbench tools (43). The increased values of strength and iterations during smoothing are due to the lack of a mid-thickness surface, which requires further corrections to properly space out the sulci.

In the walrus brain, although there was a visible contrast between white and grey matter, the presence of a physical artefact on the brain rendered the usage of ITK-SNAP and automatic segmentation impossible. Therefore, we first reconstructed both pial and white matter surfaces with Stitcher and subsequently ran the mris_volmask from FreeSurfer to create very precise segmentation masks, minimizing the effects of damaged tissue present in the MRI. Finally, we ran the default pipeline for precon_all for the final analysis. The first reconstruction *via* Stitcher was used to better visualize and understand the anatomy of damaged regions. Since the software always generates deterministic results, it allows a positive feedback loop from manual segmentation to reconstruction, then to surface analysis, and finally back to the segmentation, where small fixes can be made. As Stitcher relies solely on contours rather than image contrast, this also means the final mask is less affected by MRI artefacts and that anatomical expertise can be better incorporated into the analysis.

Irrespective of the reconstruction approach, all final cortical surfaces were converted to GIFTI surface format (.surf.gii) and subsequently processed using the same surface-based workflow for sulcal labelling and analysis.

#### 2. Supplementary figures

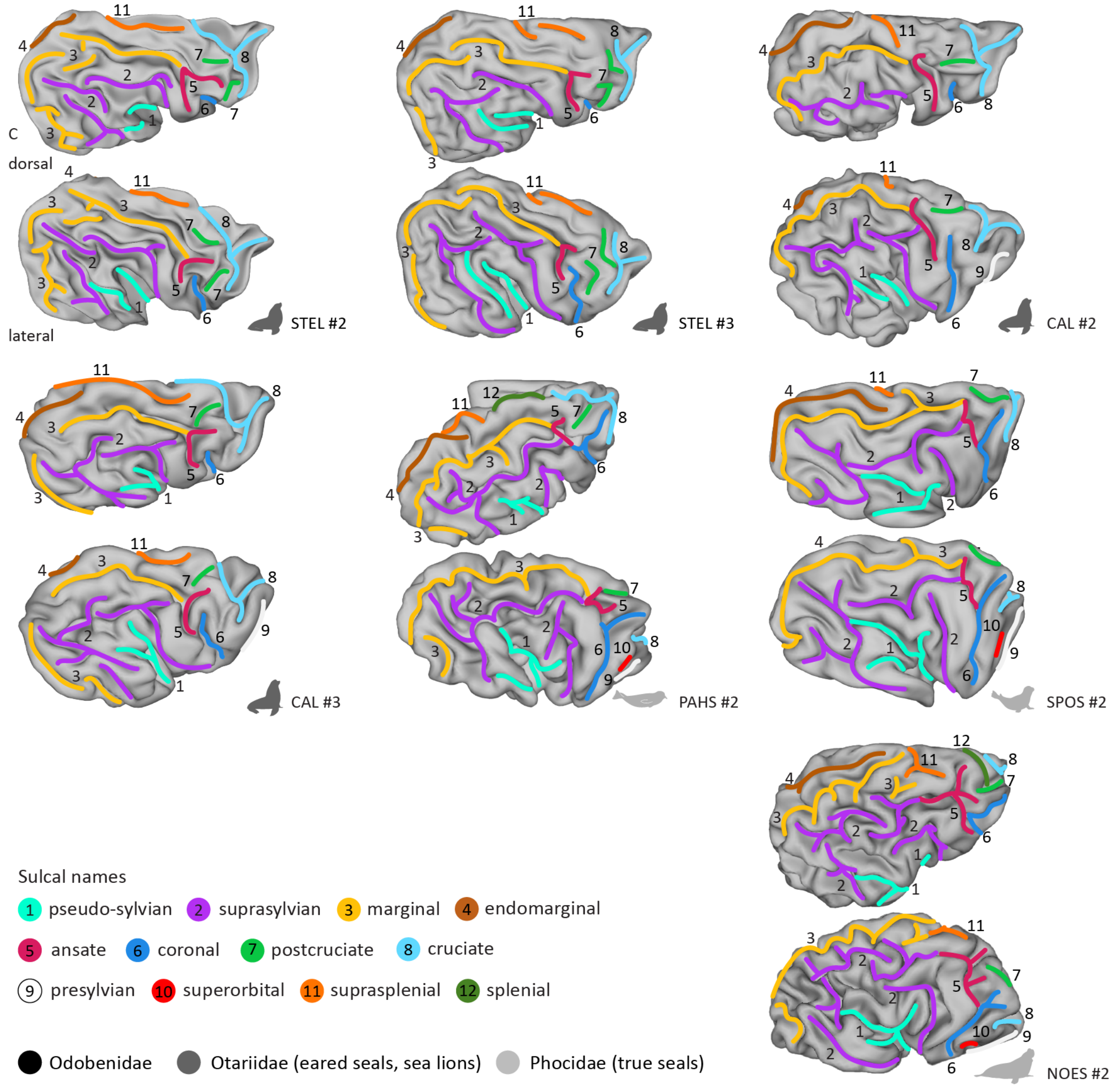


**Supplementary Figure 1. Lateral and dorsal views of additional individuals confirm sulcal patterns observed in individuals described in the main text.** Sulcal anatomy is shown for additional individuals of Steller sea lion (STEL), California sea lion (CAL), Pacific harbour seal (PAHS), spotted seal (SPOS), and northern elephant seal (NOES), illustrating relative intraspecific variation in sulcal shape, continuity, and branching. Surfaces are arranged by pinniped family, indicated by icon colours: dark grey, Otariidae; light grey, Phocidae; see **Table 1** for sample details.

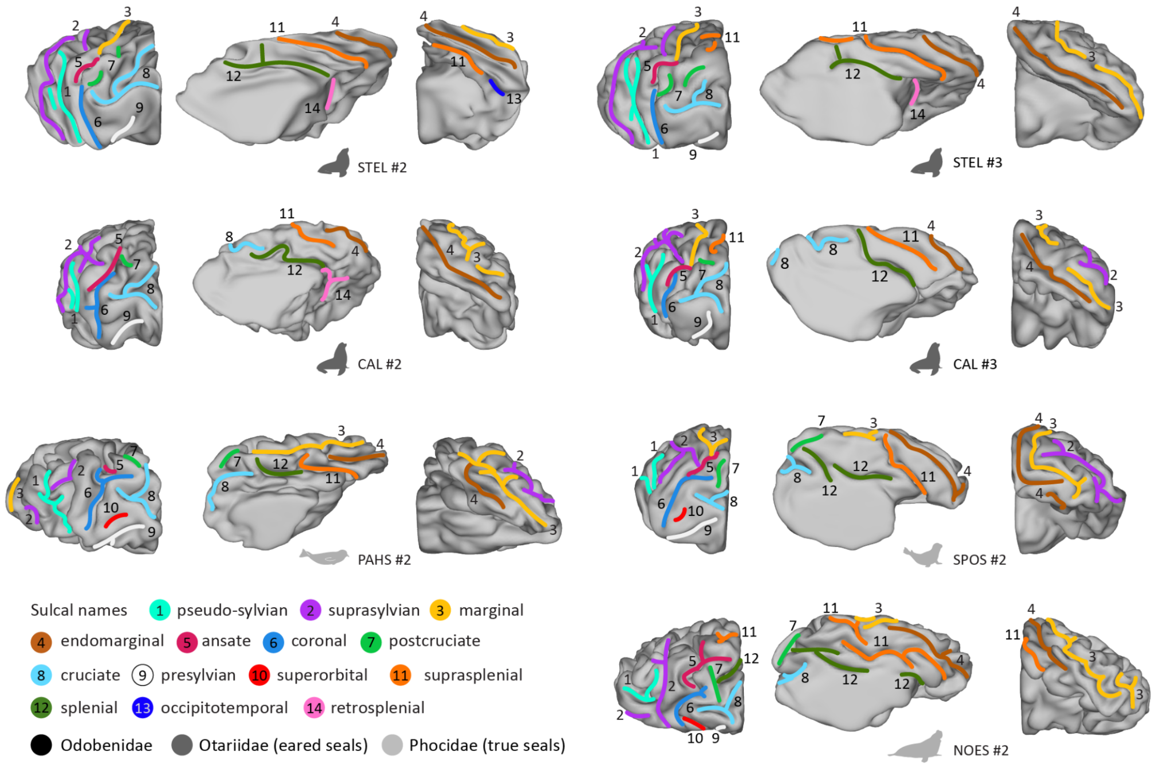


**Supplementary Figure 2. Rostral, medial, and caudal views of additional individuals confirm the major sulcal patterns described in the main sample.** Sulcal anatomy is shown for additional individuals of Steller sea lion (STEL), California sea lion (CAL), Pacific harbour seal (PAHS), spotted seal (SPOS), and northern elephant seal (NOES), illustrating relative intraspecific variation in sulcal shape, continuity, and branching, including sulci of the medial wall and their extension onto the dorsal and caudal cortical surfaces. Surfaces are arranged by pinniped family, indicated by icon colours: dark grey, Otariidae; light grey, Phocidae; see **Table 1** for sample details.

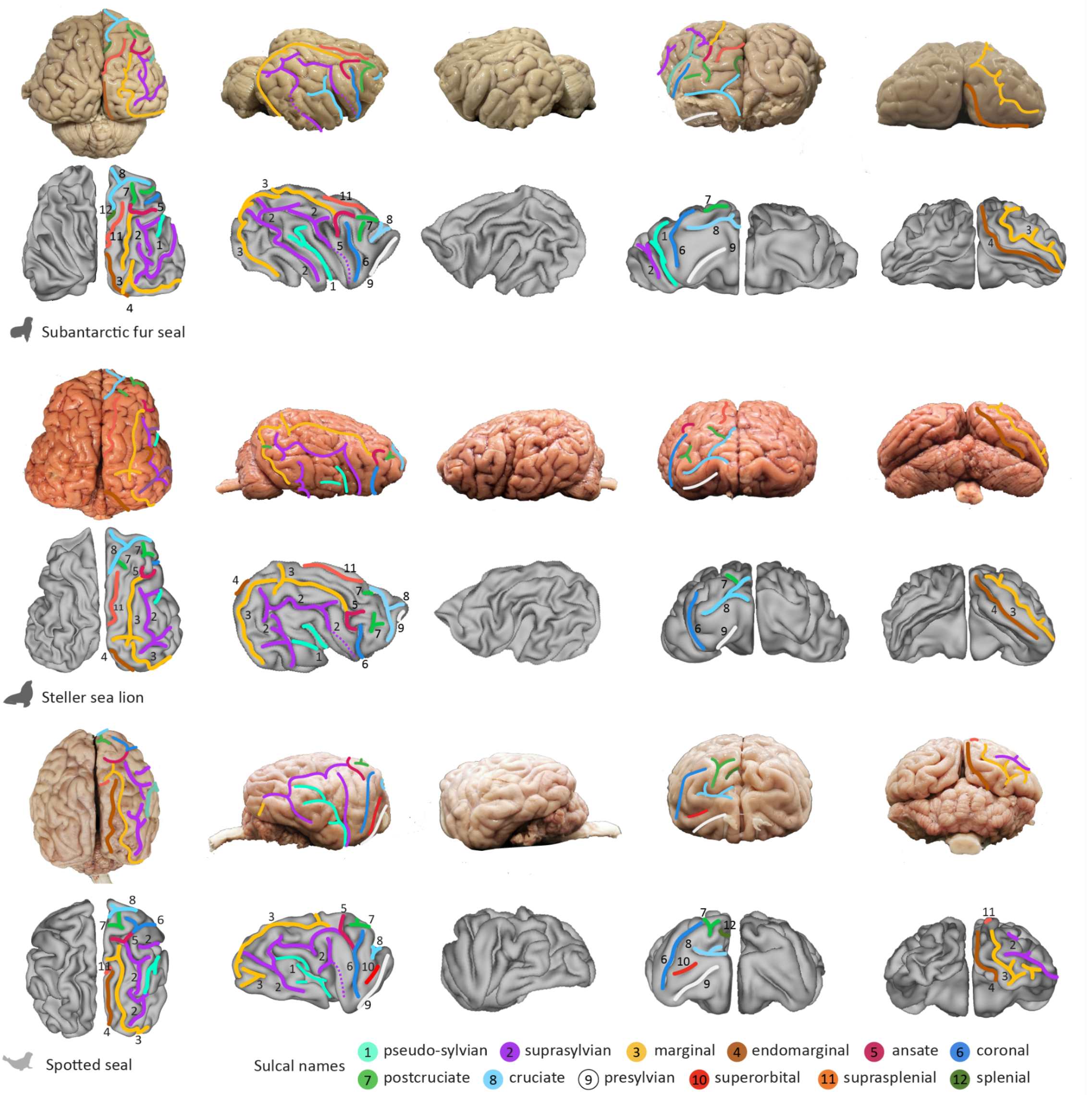


**Supplementary Figure 3. Photographs from dorsal, lateral, frontal and caudal views of three different pinniped species confirm the major sulcal patterns described in the main sample.** Sulcal anatomy is shown in post-mortem species (photographs) and corresponding digital reconstructions for the Subantarctic fur seal (SAFS), the Steller sea lion (STEL), and the Spotted seal (SPOS), illustrating consistency between modalities, highlighting via dotted lines the trajectory of the pseudosylvian and rostral suprasylvian sulci where they lie hidden beneath adjacent gyri.

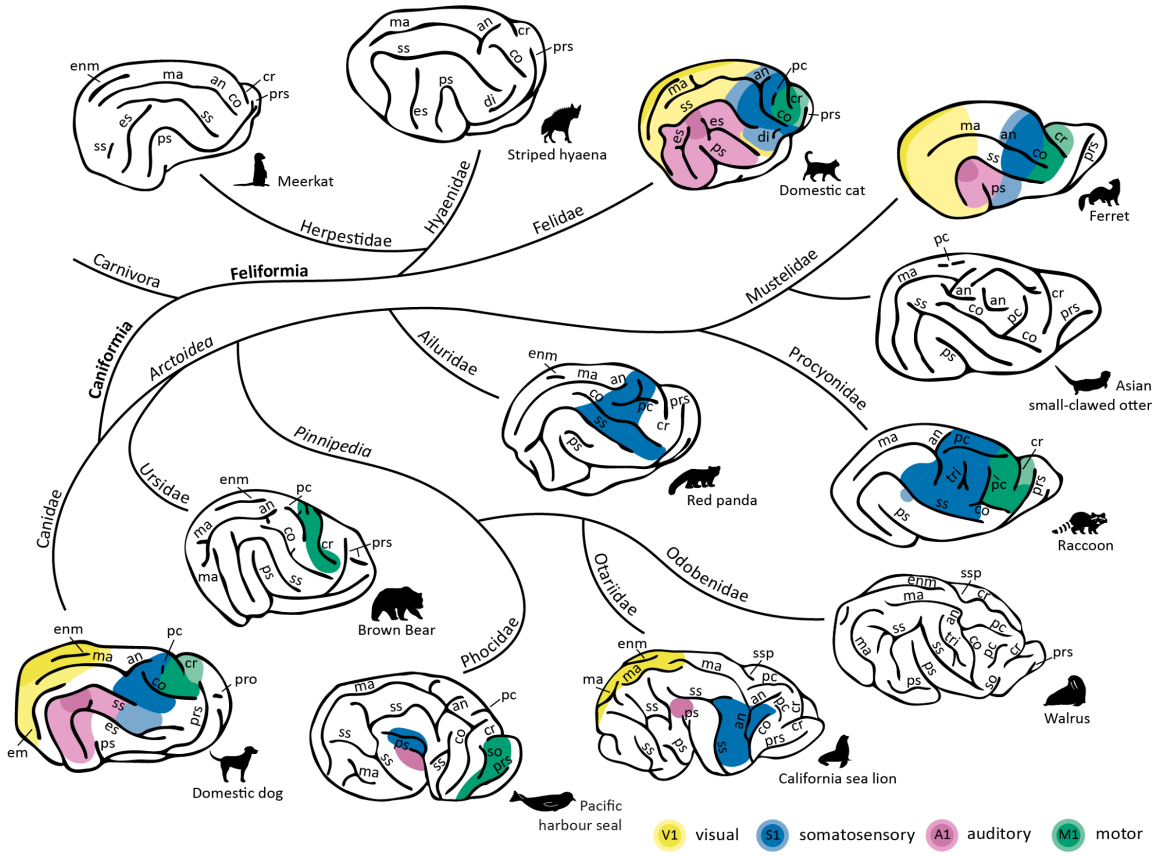


**Figure 6. Carnivoran brain diversity illustrated by representatives of eleven families**. Sulcal anatomy and functional cortical territories across representatives of eleven carnivoran families. Schematic lateral views show major neocortical sulci and the approximate locations of primary visual (V1), somatosensory (S1), auditory (A1), and motor (M1) cortical territories, combining pinnipeds examined here with fissipeds from [21]. Functional territories are shown where supported by previous anatomical, electrophysiological, or neuroimaging studies; uncoloured territories indicate unavailable or unrepresented functional information. an, ansate; co, coronal; cr, cruciate; di, diagonal; em, ectomarginal; enm, endomarginal; es, ectosylvian; ma, marginal; pc, postcruciate; pro, proreal; prs, presylvian; ps, pseudosylvian; so, superorbital; ss, suprasylvian; ssp, suprasplenial; tri, triradiate.
